# Human spermatogonial stem cells retain states with a foetal-like signature

**DOI:** 10.1101/2024.03.18.585628

**Authors:** Stephen J. Bush, Rafail Nikola, Seungmin Han, Shinnosuke Suzuki, Shosei Yoshida, Benjamin D. Simons, Anne Goriely

## Abstract

Spermatogenesis involves a complex process of cellular differentiation maintained by spermatogonial stem cells (SSCs). Being critical to male reproduction, it is generally assumed that spermatogenesis starts and ends in equivalent transcriptional states in related species. Based on single-cell gene expression profiling it has been proposed that undifferentiated human spermatogonia can be sub-classified into four heterogenous subtypes, termed states 0, 0A, 0B, and 1. To increase the resolution of the undifferentiated compartment and trace the origin of the spermatogenic trajectory, we re-analysed the single-cell (sc)RNA-seq libraries of 34 post-pubescent human testes to generate an integrated atlas of germ cell differentiation. We then used this atlas to perform comparative analyses of the putative SSC transcriptome both across human development (using 28 foetal and pre-pubertal scRNA-seq libraries) and across species (including data from sheep, pig, buffalo, rhesus and cynomolgus macaque, rat and mouse).

Alongside its detailed characterisation, we show that the transcriptional heterogeneity of the undifferentiated spermatogonial cell compartment varies not only between species but across development. Our findings associate ‘state 0B’ with a suppressive transcriptomic program that, in adult humans, acts to functionally oppose proliferation and maintain cells in a ready-to-react state. Consistent with this conclusion, we show that human foetal germ cells – which are mitotically arrested – can be characterised solely as state 0B. While germ cells with a state 0B signature are also present in foetal mouse (and are likely conserved at this stage throughout mammals), they are not maintained into adulthood. We conjecture that in rodents, the foetal-like state 0B differentiates at birth into the renewing SSC population, whereas in humans it is maintained as a reserve population, supporting testicular homeostasis over a longer reproductive life while reducing mutagenic load. Together, these results suggest that SSCs adopt differing evolutionary strategies across species to ensure fertility and genome integrity over vastly differing life histories and reproductive timeframes.

## Introduction

To maintain sperm production over the reproductive lifespan, spermatogonial stem cells (SSCs) and their progenies regularly undergo multiple rounds of mitotic division. Despite this constant demand, germline mutation rates remain significantly lower than those observed in somatic cells ^1^. Although the underlying protection strategies remain in question, it is thought that they relate to DNA replication fidelity and damage ^2^, and that the regulation of SSC turnover may play a fundamental role ^3,4^. To define the transcriptional signature of germ cell states and their potential conservation across species, emphasis has been placed on the single-cell profiling of whole testes. However, the scarcity and variability of the undifferentiated spermatogonial population and the inherent stochasticity of single-cell transcriptomics data has confounded the detailed characterisation of the stem cell compartment ^5,6^.

Historically, models of human spermatogenesis date back to the 1960s and posit the existence of two subtypes of type A spermatogonia, A_dark_ and A_pale._, both with undifferentiated nuclear morphology but differential staining intensity with hematoxylin ^7,8^. Within this model, only A_pale_ function as actively-dividing stem cells ^9^ while A_dark_ constitute a quiescent population that acts as a regenerative reserve should, for example, the A_pale_ pool be lost to gonadotoxic insult. However, in recent years this ‘linear’ model has been challenged. Through advances in single-cell transcriptomics, successive studies based on the profiling of whole human testes ^10,11^ have found evidence of multiple discrete states of type A spermatogonia, none of which unambiguously correlate with an A_dark_ or A_pale_ phenotype ^12^. Furthermore, in a previous study applying hematoxylin-staining to human seminiferous tubules, many cells were found to have an intermediate morphology, simultaneously resembling both A_pale_ (by staining lightly with hematoxylin) and A_dark_ (possessing a non-staining nuclear rarefaction zone) ^13^. There is now an emerging consensus that the undifferentiated fraction of human spermatogonia can be sub-classified into at least four subtypes (reviewed in ^14^ and based largely on the work of two research groups ^15,16^), each stably maintained throughout reproductive life: state 0 (*EGR4*+ *PIWIL4*+ *TSPAN33*+), state 0A (*FGFR3+*), state 0B (*NANOS2*+), and state 1 (*GFRA1*+ *NANOS3*+). Among these, state 0 is considered the most undifferentiated, standing at the apex of the differentiation trajectory. Contrary to the strictly hierarchical A_dark_/A_pale_ model, the existence of at least four transcriptional states suggests a more complex and dynamic SSC regulation. In common with mouse spermatogenesis ^4,17^, it has been proposed that to confer resilience upon the SSC pool and maintain homeostasis over the long reproductive lifespan, humans rely on a pattern of stochastic SSC renewal in which stem cell loss through differentiation is compensated by the duplication of neighbours, leading to a clonal dynamics characterised by neutral drift. Such behaviour would also be compatible with the phenomenon of selfish spermatogonial selection, a process whereby certain pathogenic *de novo* mutations are transmitted to progeny at an apparent mutation rate up to 1000-fold higher than background, because they provide a selective advantage to mutant SSCs that clonally expand as men age ^18,19^.

Despite advances in single-cell transcriptomics, few human testes datasets are available and, accordingly, the molecular identity of the undifferentiated compartments and their regulation remain in question. Specifically, it is still unclear as to which transcriptional programs promote SSC development, how SSCs are regulated to maintain lifelong spermatogenesis, and the extent to which these programs are conserved across species. As spermatogenesis is conserved in mammals, it is also generally assumed that germ cell differentiation starts and ends in equivalent transcriptomic states in related species (although it may not proceed at the same rate through intermediary states)^20^. For instance, a recent study used single-nucleus RNA-sequencing (snRNA-seq) to profile the germline of ten mammals and a bird, characterising spermatogenesis as a common and continuous path of cellular proliferation and differentiation that in each species originated in an analogous, but undissected, SSC cluster ^21^. In general, SSCs are underexplored primarily due to the limited number of spermatogonia within the testis ^6^ and the difficulty in procuring samples.

Here, we hypothesise that an integrated single-cell atlas of the set of publicly-available adult human testes samples (comprising data from 9 different studies ^10,15,16,20,22–26^) would increase the resolution at which we view the molecular identity of the undifferentiated compartment and the origin and progression of the spermatogenic trajectory. Combined with a comparative analysis of spermatogonia across human development (using embryonic, foetal and both pre– and peri-pubertal samples from 7 different studies ^10,16,23,25,27–29^) and against two primate (rhesus ^5,20^ and cynomolgus macaque ^30^), and five non-primate mammalian species (mouse ^31–34^, rat ^35^, pig ^36^, sheep ^37^, and buffalo ^38^), here we aim to characterise the heterogeneity of undifferentiated human spermatogonia and provide insight into the nature of the putative SSC populations. In doing so, we propose new hypotheses about the mechanisms of SSC regulation. Specifically, we show that the transcriptional heterogeneity of the undifferentiated compartment varies not only across species but through development. Further, based on the projection of a time-series of human single-cell datasets, we suggest a role for the recently-characterised state 0B as a suppressive transcriptomic program that in adults acts to functionally oppose proliferation and maintain SSCs in a mitotically poised or ‘ready-to-react’ state. Supporting this conclusion, we show that foetal germ cells – which are mitotically arrested – share the transcriptional signature of state 0B. Finally, we conjecture that the relative presence and absence of analogous transcriptomic states suggests that the mechanisms of adult spermatogenesis may differ across species. We argue that this divergence reflects differing evolutionary strategies to balance reproductive success with the control of germline mutation rate, ensuring the transmission of high-quality genetic material despite vastly differing life histories and reproductive timeframes.

## Results

### A scRNA-seq atlas of adult human spermatogonial stem cells

To generate an integrated single-cell atlas of human spermatogonial stem cells and their progenies, we obtained 34 post-pubescent (14-66 years) scRNA-seq testes libraries, representing data from 29 individuals across 9 previous studies ^10,15,16,20,22–26^ (detailed in **Supplementary Tables 1 and 2**). Samples were reprocessed using a common Kallisto/Bustools ^39^ and Seurat ^40^ workflow with conservative parameters (see Materials and Methods) to generate an integrated atlas of 60,427 testicular cells (**Figure 1**), each expressing on average 2877 genes/cell (**Supplementary Table 3**). Using an unsupervised clustering approach, we identified 10 distinct cell clusters, which we annotated using both established biomarkers for the major somatic and germ cell types (**Figure 1**; biomarkers sourced from previous publications ^10,16,25,41–44^ and detailed in **Supplementary Table 4**) and an all-against-all differential gene expression analysis (**Supplementary Table 5**). Following batch correction (see Materials and Methods), the atlas showed no residual batch effects on the basis of sample accession, study of origin, donor age or cell cycle phase (**Supplementary** Figure 1), nor an overtly disproportionate contribution made by any individual sample on the basis of number of genes, mitochondrial genes, or UMIs per cell (**Supplementary Figure 2**). The major cell types included undifferentiated spermatogonia (*UTF1*+ *EGR4*+), differentiating spermatogonia (*KIT*+ *MKI67*+), spermatocytes (*SYCP2*+ *SPATA8*+), early spermatids (*SIRT2*+ *TEX29*+), late spermatids (*PRM1*+ *PRM2*+), Sertoli cells (*AMH*+ *WT1*+), endothelial cells (*CD34*+), peritubular myoid cells (*ACTA2*+ *MYH11*+), Leydig cells (*DLK1*+ *INHBA*+) and macrophages (*CD163*+ *CSF1R*+) (**Figure 1** and **Supplementary Table 5**). Sertoli cells comprised the smallest cluster, consistent with the fact that they are especially sensitive to digestion and size selection, essential steps within many of the respective scRNA-seq protocols ^45^.

**Figure 1.**
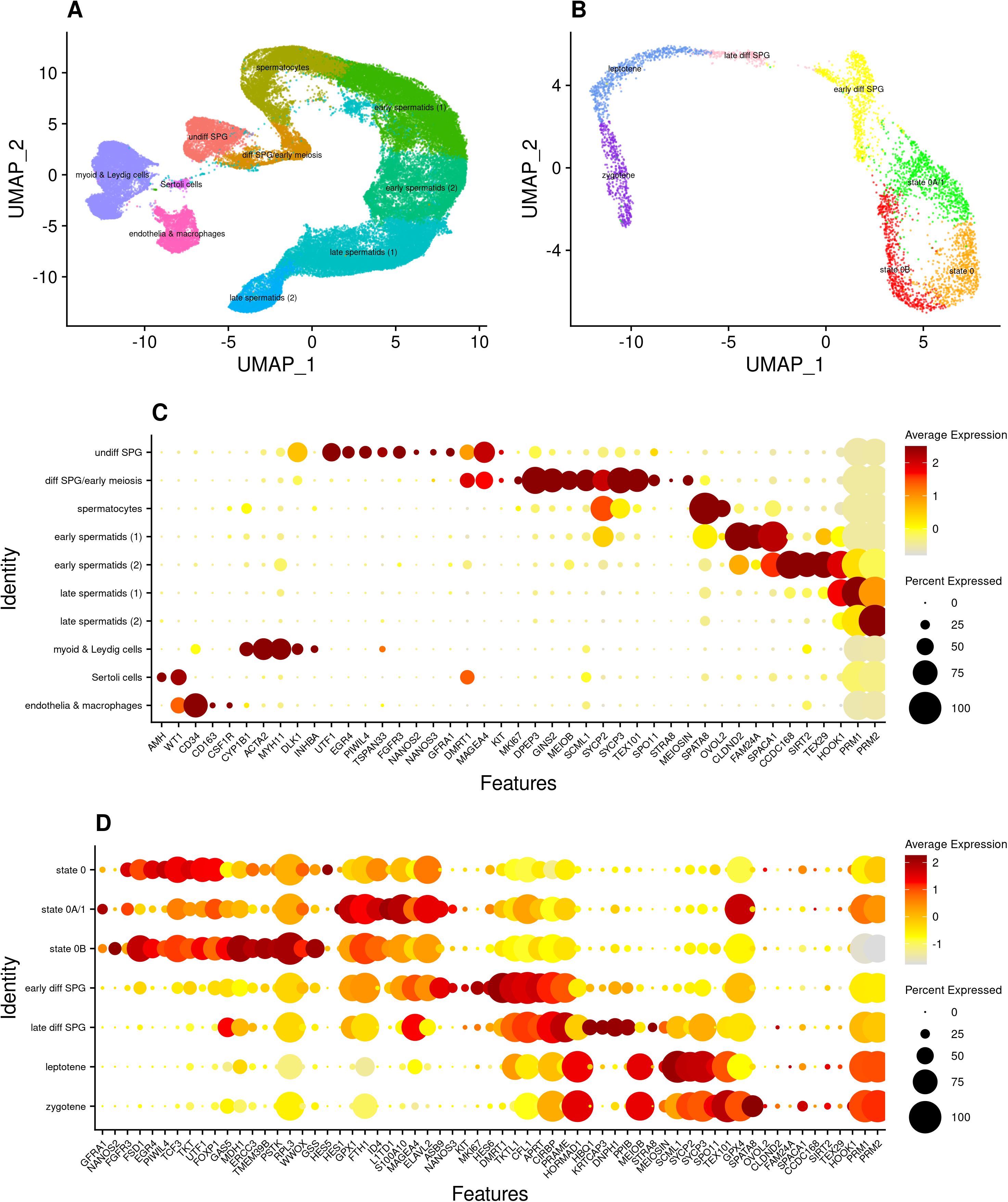
Single-cell expression atlases of. (A) the adult human testis (n = 60,427 cells), (B) a re-clustered subset comprising those cells annotated ‘undifferentiated spermatogonia’ and ‘differentiating spermatogonia’ (n = 4447 cells), hereafter the ‘SPG atlas’. Dot plots showing the relative expression of discriminative biomarkers for each cell type or state are given for both (C) the testis atlas, and (D) the SPG atlas. Note that the subset of cells comprising the SPG atlas follows a developmental trajectory intentionally broken at the onset of the meiotic program and that owing to discrete cluster boundaries being imposed upon a continuum of cells, the ‘differentiating spermatogonia’ cluster evidently comprises some primary spermatocytes (which appear as the ‘leptotene’ and ‘zygotene’ clusters of the SPG atlas). Contents of these testis and SPG atlases, showing expression level per gene per cell cluster, are available as **Supplementary Tables 5 and 7**, respectively. Variants of the SPG atlas at different clustering resolutions are shown in **Supplementary Figure 8** and presented in **Supplementary Tables 6 and 8**.

As our primary focus was to resolve germ cell rather than somatic cell types, we empirically selected the minimum clustering resolution necessary to relate the germ cell clusters to established cell types, rather than risk over-clustering and segregating cells on the basis of non-biological variation. While this approach maximises the inclusion of undifferentiated spermatogonia in a single cluster (for subsequent refinement, discussed below), this approach may under-cluster some of the somatic cell types and mask their underlying biological structure. To that end, myoid and Leydig cells were aggregated into one supercluster, as were endothelial cells and macrophages (**Figure 1**). As an independent validation of data integrity, and reinforcing the annotation of the two spermatogonial clusters (‘undiff SPG’ and ‘diff SPG’), we found that of a list of 43 genes previously detected at the protein level only in human spermatogonia, 23 (54%) were found to be differentially expressed only in either of these two clusters and no other germ cell cluster (specifically, *CBL, CD9, CHEK2, DMRT1, DMRTB1, ELAVL2, EPCAM, EXOSC10, FGFR3, FMR1, ID4, ITGA6, ITGB1*, *MAGEA4, PASD1, PHF13, POU2F2, SPOCD1, SSX3, TRAPPC6A, UCHL1, UTF1*, and *ZBTB16*) ^12^.

These two clusters of (predominantly) pre-meiotic germ cells were extracted and re-integrated to create an adult ‘SPG atlas’ of 4447 cells (**Figure 1**), which we also re-clustered using an unsupervised approach. As with the whole-testis dataset, we found no evidence of batch effects (**Supplementary Figure 3**) nor the disproportionate contribution by individual samples (**Supplementary Figure 4**). However, as spermatogonia constitute a small proportion of the total number of testicular cells and their meaningful integration could not be made for samples with < 100 cells (as integration requires the identification of comparable cell populations across multiple samples), the atlas was restricted to data from 9 (26%) of the original 34 libraries, representing 5 of the 9 original studies (more specifically, data from refs. ^10,22,25,26^ were excluded). Nevertheless, each contributing sample and each study made an approximately proportionate contribution of UMIs to each of the spermatogonial clusters, such that downstream analyses were unlikely to be influenced by residual sample– or study-specific effects (**Supplementary Figure 5**). Reciprocal projection of the SPG atlas onto the whole-testis atlas, and of the ‘undiff SPG’ and ‘diff SPG’ clusters of the whole-testis atlas onto the SPG atlas, further confirmed that the two atlases shared the same cell populations (**Supplementary Figure 6**). The expression of key marker genes is shown in **Supplementary Figure 7** and used to inform annotation, discussed below.

To further develop our analysis, we then sought to sub-type the undifferentiated compartment (depicted as the ring-like structure in **Figure 1B**), noting that the number of cell clusters within this population is algorithmically derived, a function of Seurat’s ‘cluster resolution’ parameter. We first used Clustree ^46^ to visualize the effect of different resolutions, finding the number of sub-states varied stepwise from two to four at resolutions 0.3, 1.1 and 1.2, respectively (see **Supplementary Figures 8 and 9**, with the respective datasets available as **Supplementary Tables 6 to 8**). For subsequent analysis, we clustered the atlas at resolution 1.1 on the basis that it split the ring into three distinct clusters that could each be related to the existing literature (**Figure 1**), discussed more fully below.

At this resolution, the SPG atlas comprised 7 distinct clusters, representing both undifferentiated states and distinct differentiating cell types, which we annotated on the basis of established biomarkers (**Supplementary Table 4**) and global differential gene expression analysis (**Supplementary Table 8**). To attach meaningful names to each cluster, we noted that three recent studies (Di Persio ^15^, Sohni ^16^ and Guo ^10^) reinforced each other by defining analogous undifferentiated cell states, albeit with varying nomenclature (**Supplementary Table 4**). We adopted the terminology of Guo ^10^ and Di Persio ^15^, and annotated the three undifferentiated cell clusters as “state 0” (*EGR4*+ *PIWIL4*+), a combined “state 0A/1” (*FGFR3*+ *GFRA1*+), and “state 0B” (*NANOS2*+), respectively, with the remaining four clusters of differentiating cells annotated as “early differentiating spermatogonia” (early diff SPG; *DMRT1*+ *KIT*+ *MKI67*+), “late differentiating spermatogonia” (late diff SPG; *STRA8*+), “leptotene” (*SYCP3*+) and “zygotene” (*SPATA8*+).

Note that although the same dataset produce two undifferentiated cell clusters from resolutions 0.3 to 1.0 and four from 1.2 to 2.0 (we did not continue clustering at higher levels), the cells on the right-hand side of the ring formed two stable clusters from resolution 1.1 onwards, and those on the left-hand side formed either one (resolution 1.1) or two (resolution 1.2 and higher) clusters, respectively (discussed in more detail below). Nevertheless, without foreknowledge of the biological significance of these putative undifferentiated states, it is difficult to determine whether the cells have been under– or over-clustered at a given resolution ^47^. Under-clustering masks biological structure by assigning cells to clusters that are too broad, whereas over-clustering segregates cells on the basis of non-biological variation. In this respect, resolution 1.1 may be considered ‘optimally’ clustered as there were no apparent noisy or artefactual populations, and each of its clusters could be related to known cell states without defining new ones (i.e., the two subtypes of state 0B, which form at resolution 1.2 and higher, discussed below). However, we note no clear distinction between state 0A and state 1, which were considered separate *FGFR3*+ and *GFRA1*+ *NANOS3*+ populations by ^15^ (in our SPG atlas, *FGFR3* and *GFRA1* are differentially expressed in the same undiff SPG cluster whilst *NANOS3*+ is differentially expressed in ‘early diff SPG’; **Supplementary Table 7**). Regardless, it is important not to lose sight of the fact that these are discrete states imposed on what may in reality be more of a continuum, with some studies suggesting a gradual transition between states rather than binary on/off programs ^10,15,24,44^. Indeed, consistent with this, there were very strong correlations between the expression profiles of each of the three undifferentiated cell clusters, which primarily differ not in the identity of the genes they express, but in their relative levels of expression (**Supplementary Figure 10**).

Of the 4447 cells in the germ cell atlas, 2229 (50%) were undifferentiated (i.e. the total number of cells annotated as states 0, 0A/1 and 0B, collectively forming the **Figure 1B** ring). Note that in their respective UMAPs, the studies by Sohni ^16^ and Di Persio ^15^ also identified this ring-like structure, and noted the differential enrichment of *FGFR3* and *NANOS2* on opposite sides (as also seen in **Supplementary Figure 7**). However, neither study speculated in much detail as to its origin and biological significance. Here, using the larger integrated datasets, we aimed to refine and expand upon these findings.

### Sub-populations of undifferentiated spermatogonia with differing metabolic profiles

At the chosen level of clustering (resolution 1.1), the three subpopulations of undifferentiated spermatagonia segregated into states at the bottom (state 0), right (state 0A/1), and left (state 0B) of the ring-like structure shown in **Figure 1B**. The cell population immediately above the ring constitute differentiating spermatogonia, with the apparent gap between this cluster and the onset of the meiotic program likely reflecting both the conservative thresholds employed to generate the atlas and the capture of few cells at this stage of differentiation.

To characterize the nature of the undifferentiated subpopulations, we first applied RNA velocity analysis, a method of inferring cell fate trajectories from the relative abundance of nascent (unspliced) and mature (spliced) mRNA ^48^, illustrated in **Supplementary Figure 11**. For the cells within the ring, we made three observations. First, cells in state 0 appeared to be, for the most part, transcriptionally ‘undirected’, consistent with their potential identity as a ‘reserve’ population and the starting point (i.e., apex) of the spermatogonial trajectory. Second, vectors aggregate at the very top of the ring, in state 0A/1, with arrows oriented upwards from both state 0A and state 0B, and downwards from ‘early diff SPG’, and third, within state 0B, there are two sets of streamlines, one pointing towards state 0A/1 and another towards state 0 (note that at resolution 1.2, the state 0B cluster splits into two on this basis; see **Supplementary Figure 9**). Velocity plots are characteristically challenging to interpret although taken together these observations suggest that undifferentiated spermatogonia could be transcriptionally orientated both towards and away from state 0, such that the spermatogonial trajectory may not proceed linearly from a state 0 ‘start’. Rather, the transcriptomic programs of 0A and 0B may be interpreted as, in part, directed towards and away from a commitment to differentiation, respectively.

To explore this concept further, we performed GO and KEGG enrichment analysis on the set of genes differentially expressed (DE) in each cluster, using both all-against-all and pairwise comparisons (**Supplementary Tables 10 to 12**). Consistent with the energetic expense of differentiation, cells in state 0A/1 had a metabolic profile shifted towards oxidative phosphorylation (OXPHOS). GO terms enriched among genes DE in state 0A/1, relative to the other six clusters in the atlas (i.e., including undifferentiated and differentiating spermatogonia, and cells at the onset of meiosis), included ‘aerobic respiration’, ‘ATP metabolic process’ and ‘mitochondrial respiratory chain complex assembly’ (**Supplementary Table 10**). Furthermore, a pairwise analysis of genes differentially expressed in state 0A/1 relative only to either states 0 or 0B showed enrichment for essentially the same terms, including ‘mitochondrial ATP synthesis coupled electron transport’ and ‘aerobic electron transport chain’ (**Supplementary Table 11**). The implication is that spermatogonia in the other undifferentiated states have a relatively hypoxic metabolic profile, with cells in state 0A having transitioned away from this. Also consistent with this, DE genes in state 0A/1 relative to state 0 were enriched for the term ‘regulation of transcription from RNA polymerase II promoter in response to hypoxia’ (**Supplementary Table 11**). We note, however, that there was no direct evidence of enriched GO or KEGG terms related to (for example) glucose metabolism and that a condition of relative hypoxia in states 0 and 0B is inferred. Conversely, however, we found that both *ECSIT*, a regulator of the balance between mitochondrial respiration and glycolysis ^49^, and *PGLS*, a component of the pentose phosphate pathway (PPP), were differentially upregulated in state 0B (**Supplementary Table 7**). The latter is of interest because PPP enzymes are downregulated in hypoxic conditions, facilitating the shift in glucose metabolism towards direct glycolysis ^50^ (see also **Supplementary Text** for additional evidence of glycolysis in state 0, using an orthogonal dataset ^38^). As state 0 appears less inclined towards OXPHOS (and thereby differentiation), we characterized this cluster in further detail.

### The ‘state 0’ or ‘SSC reserve’ cluster has little transcriptomic evidence of G_0_ quiescence

Quiescence (defined as reversible growth arrest) is a common, but not requisite, feature of many stem cell populations ^51^, including neural ^52^ and hematopoietic ^53^, which reside in hypoxic niches and rely on glycolytic metabolism. In the classical model of spermatogenesis, A_dark_ are thought to constitute a quiescent SSC reserve, replenishing the pool of actively cycling SSCs (i.e. those associated with state 0A/1, or A_pale_ in the classical model). However, on the basis of the single-cell atlas, and irrespective of any inference of hypoxia, there is no compelling evidence that as a whole ‘state 0’ is quiescent. To draw this conclusion, we considered cell cycle scoring, GO/KEGG enrichment in state 0, and the relative expression of established G_0_-markers in better-characterised stem cell systems.

First, based on their transcriptional signature, we found that the majority of undifferentiated spermatogonia were predicted to be positioned in either the S or G_2_/M phases of the cycle, i.e., actively replicating DNA, with very few cells in G_1_ (and, by implication, G_0_). Of the 725 cells in the state 0 cluster, only 90 (12%) were scored as G_1_ (**Supplementary Table 13**). Similar proportions have been reported in mouse germ cells, in which 3% of undifferentiated spermatogonia were predicted to be in G_1_ phase and the remainder actively dividing (43% in S phase and 55% in G_2_/M)^54^. Second, GO term enrichment analysis indicated a relatively high degree of cellular activity. The set of genes differentially expressed in state 0 were enriched for ‘cytoplasmic translation’, with 40% of the 159 genes annotated with this term differentially expressed in state 0 **(Supplementary Table 10**). Third, of a set of 101 genes, 68 of which comprised a quiescent gene signature from a previous study ^55^ using microarrays to identify genes consistently up– or downregulated in quiescent haematopoietic stem cells (HSCs), muscle stem cells and hair follicle stem cells, and 35 of which were considered either positive or negative intrinsic regulators of HSC quiescence (in either humans ^56^ or mice ^57^), only 9 were differentially expressed in state 0 and only 3 exclusively in state 0 (**Supplementary Table 14**). Similarly, there was neither differential expression of *GPRC5C*, considered a marker for a ‘highly dormant’ subpopulation of human HSCs ^58^, nor *DNMT3L* ^59^, an epigenetic regulator involved in the promotion of quiescence in postnatal mouse SSCs, the latter showing non-negligible expression in just 1.1% of undifferentiated cells (**Supplementary Table 7**). Further supporting the comparatively high transcriptional activity of state 0, enriched GO terms for genes differentially expressed in state 0 relative to state 0A/1 include ‘histone H4 acetylation’ (associated with the opening of chromatin) and ‘positive regulation of histone H3-K4 methylation’ (**Supplementary Table 11**). Of note, H3-K4 methylation mitigates transcription-replication conflicts (TRCs) during periods of high transcription by decelerating replication rate ^60^. In the context of a lower germline mutation rate, this is notable since TRCs, which arise when the transcription and replication machinery operate simultaneously on the same DNA template, are inherently mutagenic. Interestingly, there were no GO terms substantively enriched among the set of genes upregulated in state 0 relative to 0B. While there were 10 significantly enriched GO terms in this pairwise comparison, no term was supported by more than 7 differentially expressed genes; **Supplementary Table 11**). This lack of relatively distinct biological activity is suggestive of either a high degree of similarity between states 0 and 0B or with state 0 being an ‘undirected’ state to which a subset of cells in state 0B transition (as implied by the RNA velocity plot).

While cells in the state 0 cluster exist in a relatively hypoxic state, before the metabolic shift to OXPHOS that precedes cellular maturation and differentiation, our overall interpretation is that – contrary to what has been described before ^10^ – they are unlikely to be G_0_-quiescent. Nevertheless, it remains possible that a small and undistinguished subset of cells may still be quiescent. This would reconcile the observations of this cluster’s lack of direction on the velocity plot, its small number of ‘quiescent signature’ genes (n = 9; **Supplementary Table 14**), its relative enrichment for translational activity (a possible explanation for which is that it primarily contains cells captured immediately after they exited quiescence) and the findings of both a single-cell lineage-tracing study ^61^, which posits a quiescent SSC population marked by *FOXC2*, and a histological study ^62^ which posits a small subpopulation of undifferentiated spermatogonia as a quiescent reserve, A_dVac_ (‘A_dark_ with nuclear rarefaction zone’), potentially too few in number to have been clustered separately here. We cannot rule out the possibility that we do not directly detect an obviously quiescent pool of cells simply because current implementations of scRNA-seq are either not able to capture enough cells, or enough of the transcriptome of these cells, to meaningfully distinguish them. To that end, it is worth recalling that the data used to construct the atlas comprise unselected/unsorted cells, largely from biopsies.

### The ‘state 0B’ transcriptional program is characteristic of foetal germ cells, likely suppresses cell proliferation, and is retained into adulthood in humans

To gain further insight into the nature of states 0, 0A and 0B, we applied a two-step projection strategy following the time course of human development. The first step was to project cells from a given sample onto the whole-testis UMAP, thereby distinguishing germline from somatic cells. The second step was to project only these putative germ cells onto the SPG atlas. This strategy is conservative but necessary given that the germ cell atlas contains, by definition, only germ cells and that by projecting onto it any given sample (testicular biopsy), non-germ cells would inevitably also be projected. To validate this approach, we confirmed that FACS-sorted undifferentiated spermatogonial populations could be projected onto the map in their expected location. To this end, we obtained three sets of purified spermatogonia from a previous study ^24^, sorted against the gene combinations of HLA-ABC^negative^, CD49e^negative^, THY1^dim^, ITGA6+, and EpCAM^dim^ – markers that xenotransplantation studies have shown selectively enrich for human spermatogonia with colonization potential ^63^. We found that cells in each dataset projected almost entirely onto states 0 and 0B, as expected given that they express the markers used for enrichment (**Supplementary Figure 12**). Accordingly, we assessed whether each of the undifferentiated cell states were present throughout the human life cycle, by conservative projection of a time course of 62 samples, inclusive of the 34 samples used to construct the atlas, and incorporating foetal (8-23 week), pre– and peripubertal timepoints (**Supplementary Table 15**). Indeed, we were particularly interested in the relative contribution of the foetal samples to each undifferentiated cell state as primordial germ cells (PGCs) exist in a state of mitotic arrest ^29,54^ until their progressive re-entry into the cell cycle shortly after birth. Notably, we found that the (mitotically-arrested) PGCs of the foetal samples projected exclusively onto state 0B (**Figure 3A** and **Supplementary Table 15**). State 0B is typified by the expression of the conserved RNA-binding protein *NANOS2* (**Supplementary Figure 7**), a male-specific stem cell regulator that suppresses the proliferation and differentiation of mouse SSCs and is essential to their self-renewal ^64^. Supporting this function, conditional disruption of *Nanos2* in adult mouse testes has long been known to deplete the SSC pool, evidenced by the suppression of spermatogenesis and the loss of *Gfra1+* SSCs ^65^ (note that in mice *Nanos2*+ germ cells show high overlap with *Gfra1*+ cells ^64^ but that in humans *GFRA1*+ is characteristic of the ‘active’ state 0A/1). Nevertheless, while *Nanos2* is detectably expressed in the adult mouse testis, it has its strongest and most consistent peak of expression around embryonic days 13-15, when the germline is becoming established and PGCs are mitotically arrested ^54,66^ (see also https://tanglab.shinyapps.io/Mouse_Male_Germ_Cells/). That human PGCs also have a *NANOS2*+ profile supports the conservation of its prenatal function. During this period, PGCs in female and male gonads either, respectively, enter the meiotic program or mitotic quiescence ^67,68^. In males, the role of *Nanos2* is to suppress the former by targeting RNAs involved in meiosis, leading to their degradation ^69–72^.

**Figure 2.**
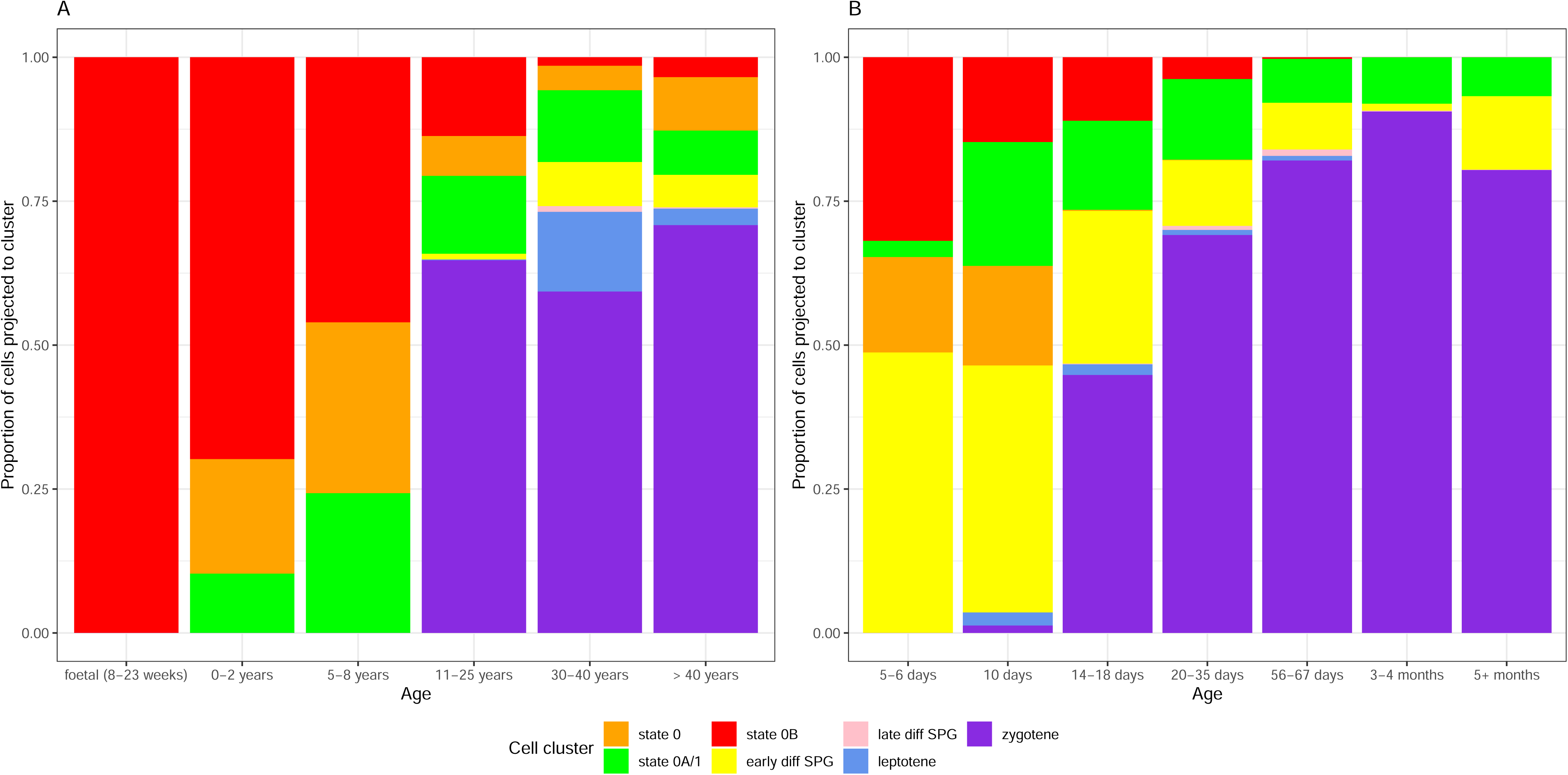
The relative abundance of undifferentiated spermatogonial states varies between humans and mice, and throughout development. Bars show the proportional number of cells projecting to each of the seven clusters of the SPG atlas (shown in Figure 1B). (A) Projection of individual human samples onto the SPG atlas (n = 62, representing 6 age groups) is consistent with the maintenance of foetal primordial germ cells in mitotic arrest, their postnatal re-entry into the cell cycle, and the onset of meiosis at puberty. That mitotically-arrested cells exclusively project to state 0B suggests a functional role for this state in suppressing cell proliferation. (B) Projection of individual mouse samples (n = 37, representing 7 age groups) shows a comparatively rapid onset of the meiotic program and the loss of both states 0 and 0B after approximately one month (i.e. sexual maturity). Details of the samples included in this analysis are given in **Supplementary Table 2**. The number of cells projected onto each cluster are given in **Supplementary Tables 15 and 17**, for human and mouse, respectively. To obtain data for this figure, the minimum projection score for both sets of projections was 0.8. Projections of individual samples are shown in the **Supplementary Text**.

**Figure 3.**
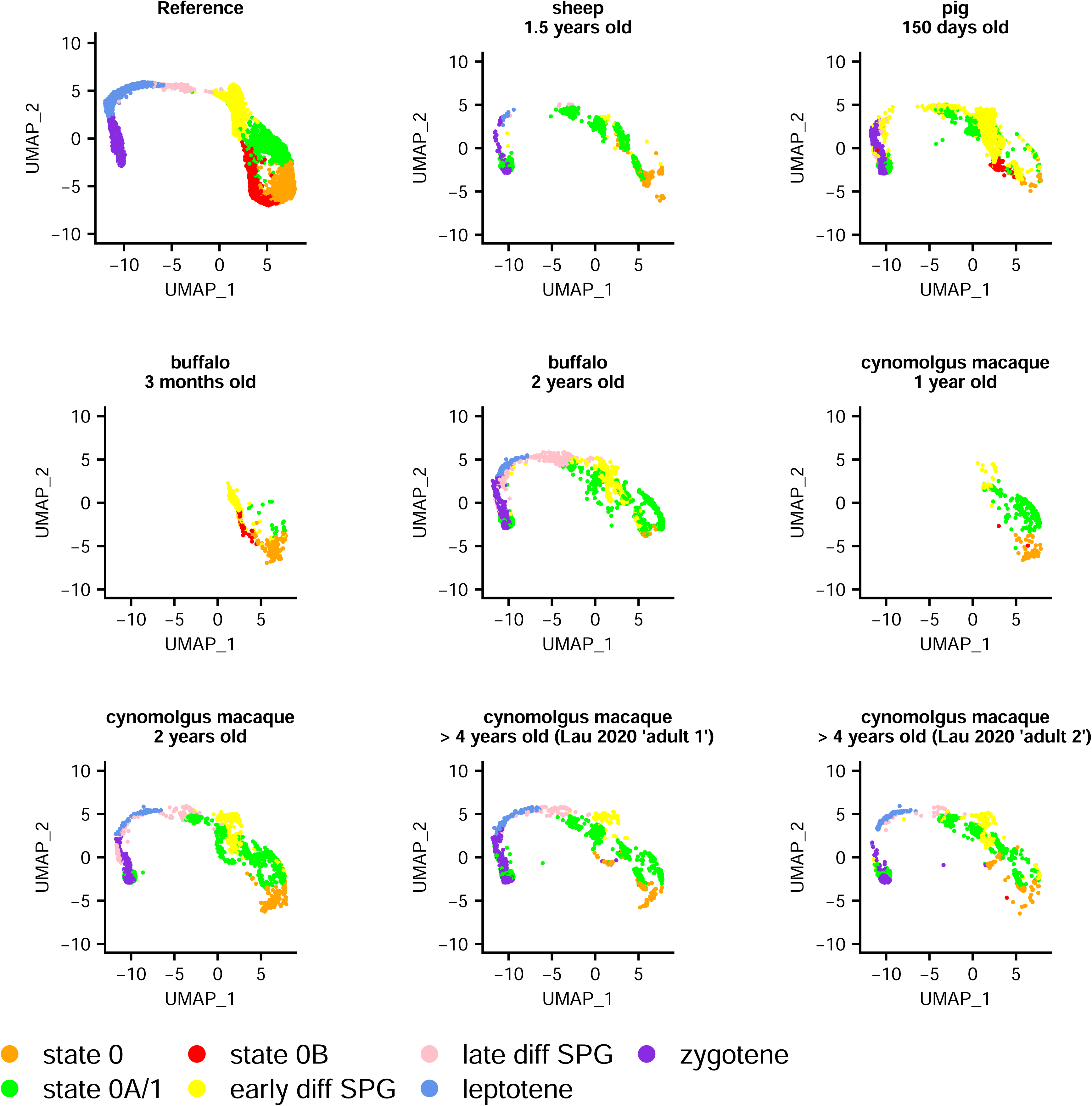
Projection of sheep, pig, buffalo and cynomologus macaque germ cells onto the SPG atlas. The 3 month-old buffalo and 1 year-old macaque samples are pre-pubertal; all others represent sexually mature individuals. These projections show that human state 0 and state 0A-like populations are consistently present in these species, but not necessarily state 0B. Details of the samples included in this analysis are given in **Supplementary Table 2**. The number of cells projected onto each cluster, for a range of minimum projection scores (0 to 0.8 at 0.2 intervals), are given in **Supplementary Table 16**, with this figure plotted with no minimum threshold required.

A more recent study that performed *Nanos2* knockout on embryonic male germ cells found that *Nanos2* expression also induces mitotic arrest by repressing the activity of mTORC1 ^73^, a key component of the mTOR signalling pathway that coordinates cell growth and metabolism in response to both environmental and intracellular stressors, including hypoxia and DNA damage ^74^. More specifically, *Nanos2* sequesters mTOR in cytoplasmic mRNPs (messenger ribonucleoproteins), a mechanism that allows for considerable flexibility as it permits transient, rapidly reversible, repression of cell growth ^75^. Conceivably, this mechanism could ‘pause’ cellular mutation independent of, or in addition to, classical (cyclin-based) mechanisms of cell cycle control. Also consistent with a ‘pausing’ or ‘suppressive’ role of state 0B, genes differentially upregulated only in state 0B (**Supplementary Table 7**) include *GAS5* (‘growth arrest-specific transcript 5’, the overexpression of which induces apoptosis and growth arrest in cancer cells) ^76^ and *MDH1* (which regulates p53-dependent cell-cycle arrest and apoptosis in response to glucose deprivation) ^77^. In addition, a pathway impact analysis (see Materials and Methods) comparing the set of genes differentially expressed in state 0B relative to state 0 found that the KEGG ‘RNA degradation’ pathway was significantly activated; by contrast, it was significantly inhibited in the ‘early diff SPG’ to ‘state 0’ comparison (**Supplementary Table 12**). Characteristics of state 0B beyond *NANOS2* upregulation are discussed further in the **Supplementary Text**.

A tantalising explanation for why proliferation would be ‘paused’ is to facilitate DNA damage repair prior to differentiation. Consistent with this, GO terms enriched among the set of genes differentially upregulated only in state 0B – which include *ERCC3*, a nucleotide excision repair gene ^78^, *TMEM39B*, which in zebrafish protects against oxidative DNA damage induced by cold stress ^79^, *PSTK*, a phosphorylase kinase with a protective role against oxidative stress ^80^, *RPL3*, involved in multiple DNA repair pathways in response to chemotherapy ^81^, and *WWOX*, which modulates the double-strand break response ^82,83^ – include ‘positive regulation of response to DNA damage stimulus’ and ‘positive regulation of apoptotic signalling pathway’ (**Supplementary Table 10**). Moreover, a pairwise ‘state 0B relative to state 0’ comparison shows enrichment for the GO term ‘positive regulation of DNA repair’ (**Supplementary Table 11**).

### Heterogeneity of undifferentiated spermatogonial states across species and development

Having established the heterogeneity of undifferentiated spermatogonial states across development, we considered to what extent they varied across species. We first projected data from five mammalian testes datasets most closely related to human (rhesus and cynomolgus macaques, plus sheep, pig, and buffalo) to determine whether they had similar undifferentiated spermatogonial states as human. In each species, cells equivalent to states 0 and 0A could be identified (**Figure 3** and **Supplementary Figures 13** and **14**). This is consistent with a shared origin of spermatogenesis in related species and a common transcriptional program underpinning SSC maturation. Curiously, however, cells projecting to state 0B belonged primarily to pre-pubertal samples, most notably for a 150-day old pig, a 3-month old buffalo, and 15-20 month old rhesus macaques (**Supplementary Table 16**), with no or a negligible number in sexually mature sheep, buffalo or macaque (**Figure 3**).

This was surprising as state 0B, in human, is distinguished largely by *NANOS2* expression alone (but see **Supplementary Text**). To rule out the possibility that the apparent absence of adult cells with a state 0B signature was an artefact of stochastic capture, we relaxed the Seurat scoring threshold used for the projections (see Materials and Methods), reducing it from a conservative 0.8 (as used in **Figure 2A**) to 0, thereby allowing the projection of any given cell to its ‘best match’ on the human UMAP. However, even with lenient parameters, state 0B cells were found to be largely absent in these non-human adults (**Figure 3**), and unsurprisingly they remained so when repeating the analysis with more conservative scoring thresholds (**Supplementary Table 16**).

Together, these results suggest that in these species state 0B plays a role in early development and that its species-specific retention into adulthood, as well as its relationship to state 0, is of particular importance to steady-state spermatogenesis. To further explore these findings, we considered at what point in development undifferentiated spermatogonial states could be ‘lost’, and why that might be. We were unable to address this with the above species as there were too few samples to reconstruct a time course of development. Accordingly, we projected an extensive time course of 37 unsorted mouse testicular samples (**Figure 2B**, **Figure 4** and **Supplementary Table 17**), starting from postnatal day (P) 5 and continuing to adulthood (detailed in **Supplementary Table 2**). In the two postnatal day (P) 5 murine testis samples, the majority of cells projected not onto the undifferentiated spermatogonial but the ‘early differentiating spermatogonia’ cluster (**Figure 4**).

**Figure 4.**
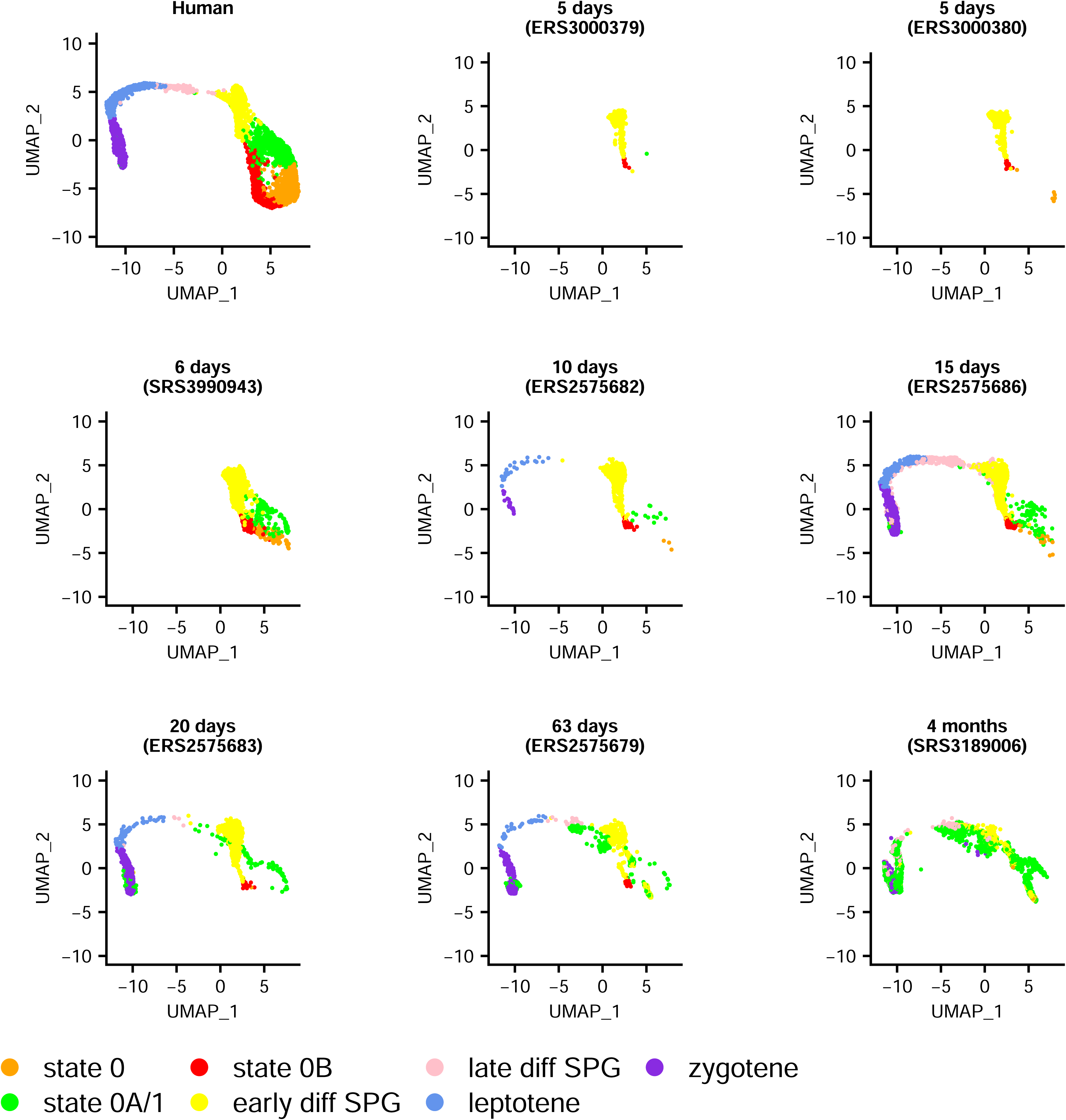
Projection of mouse germ cells onto the SPG atlas, using selected samples representing a time course from early development to adulthood. These projections show that the transcriptomic profile of mouse SSCs varies rapidly across development relative to human, with the loss within approximately one month of state 0 and 0B-like populations. Details of the samples included in this analysis are given in **Supplementary Table 2**, with the relative abundance of each cluster across the full set of 37 samples shown in Figure 2B. The full projection of every sample is given in the **Supplementary Text**. The number of cells projected onto each cluster, for a range of minimum projection scores (0 to 0.8 at 0.2 intervals), are given in **Supplementary Table 17**, with this figure plotted with no minimum threshold required.

Since in mice, progressive re-entry of PGCs into the cell cycle (after a prolonged period of mitotic arrest) commences around P1-P3 and completes by P5, this suggests that from a transcriptomic standpoint, P5 germ cells more strongly resemble those inclined to differentiate than undifferentiated spermatogonia. The P5 germ cells are also markedly different to those at P6, in which each of the human-like undifferentiated spermatogonial clusters (states 0, 0A and 0B) had been established. These observations relate to the first wave of mouse spermatogenesis ^84^, and are discussed further in the **Supplementary Text**. From P10 onwards, the meiotic program was initiated, with a concomitant loss of cells projecting to states 0 and 0B. It appears that for adult mice essentially the sole transcriptomic program of undifferentiated spermatogonia resembles human state 0A (**Figure 2B**). One possible explanation for this observation is that, as state 0A is enriched for OXPHOS metabolism and because mice, having proportionately fewer undifferentiated spermatogonia than humans ^6^, must rely on more rounds of energetically-expensive transit amplification to produce mature sperm, then in any given mouse testicular biopsy, there may simply be too few SSCs to project onto any other state. Therefore, to further confirm that cells with a human state 0 signature were absent in adult mice, we projected two sorted populations of undifferentiated germ cells (from ^85^) onto the human UMAP and obtained the same results (**Supplementary Figure 15**). Moreover, we found that when projecting additional data from 8 adult rat samples, all EpCAM+ selected and thereby enriched for germ cells, we found that as with mice they also lacked cells with states 0 or 0B signatures (**Figure 5** and **Supplementary Table 16**; EpCAM was detected in 68% of undifferentiated human spermatogonia [see **Supplementary Table 7**]). Finally, when projecting murine PGCs from embryonic day 16.5 (from ^86^) onto the human UMAP, we can see that both states 0 and 0B (but not 0A) are present (**Supplementary Figure 16**); the implication is that they differentiate at birth to form the 0A population.

**Figure 5.**
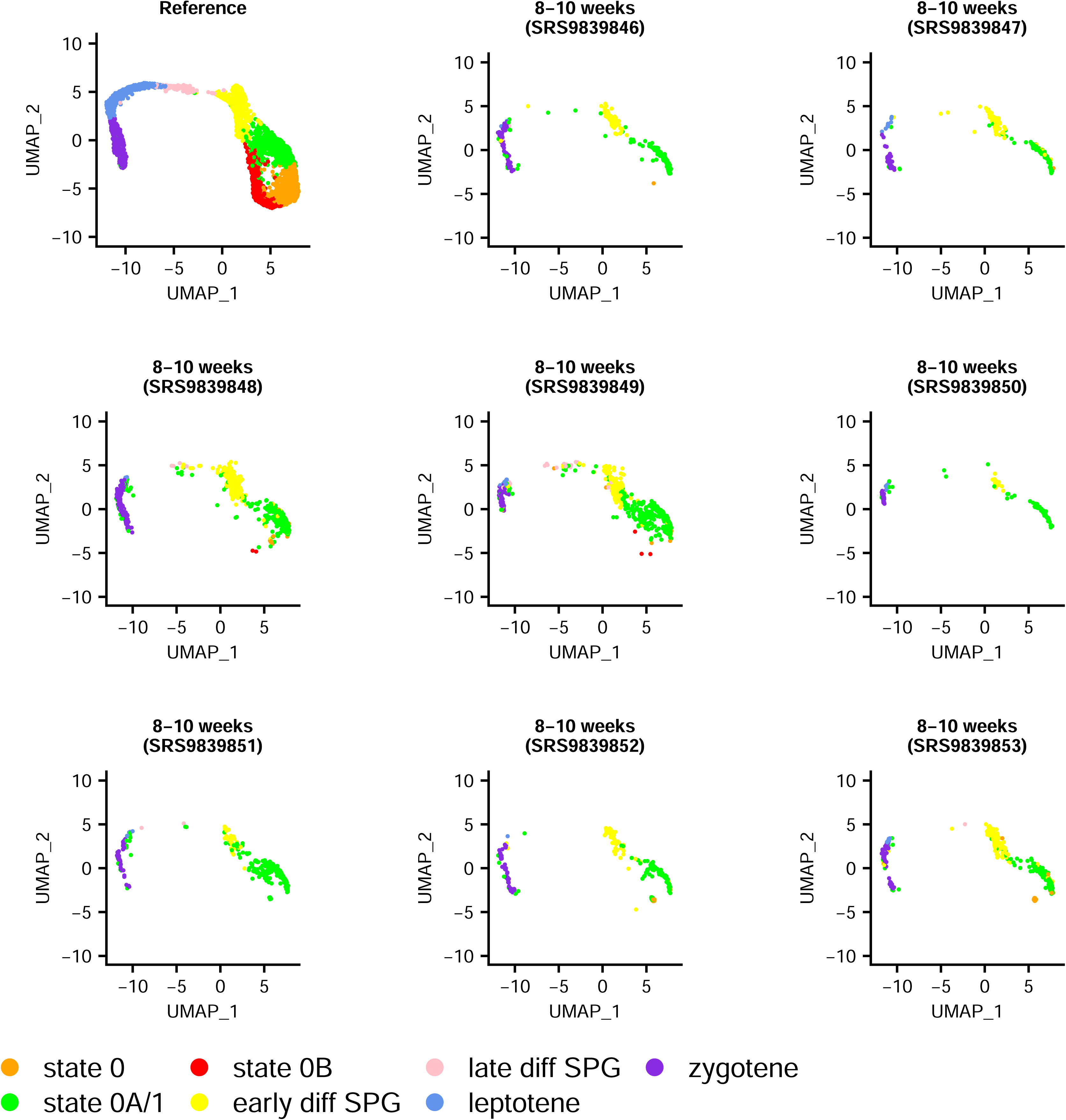
Projection of EpCAM+ selected adult rat germ cells onto the SPG atlas. These projections show that the transcriptomic profile of rat SSCs largely approximates human state 0A and that state 0 and 0B-like populations (the latter in particular) are not detectable. Details of the samples included in this analysis are given in **Supplementary Table 2**. The number of cells projected onto each cluster, for a range of minimum projection scores (0 to 0.8 at 0.2 intervals), are given in **Supplementary Table 16**, with this figure plotted with no minimum threshold required.

Overall these results suggest that, compared to the other mammals considered in this study, the spermatogenic trajectory in rodents follows a different, more developmentally mature, transcriptomic program. More broadly, these results suggest that as with many significant evolutionary steps, the history of spermatogenesis may have been shaped by heterochrony ^87^.

## Discussion

Although spermatogenesis is often viewed as a highly conserved process, our findings suggest that the origin, developmental timing and steady-state regulation of the SSC pool differs across species. While not unexpected from an evolutionary perspective ^88^, the implications of such heterogeneities have not been fully explored in previous single-cell studies, including in refs. ^20,21^, both of which posit broadly analogous transcriptional states between species across the spermatogonial trajectory – with a common origin in a seemingly homogenous ‘SSC cluster’ or ‘consensus state’. Here we have shown that contrary to this conclusion, the transcriptome of undifferentiated spermatogonia (more so than that of other, more mature, germ cells) varies greatly between species and across developmental time. In particular, we have identified a singular role of cells with a state 0B signature that is present during the pubertal development of multiple mammalian species but persists into adulthood only in human. This state, which belongs to the undifferentiated spermatogonial pool, has a mitotically poised or ready-to-react transcriptional profile, suggesting its potential role as a reserve population. The relative presence and absence of analogous undifferentiated spermatogonial states between species suggest that SSCs have adopted different strategies to maintain steady-state spermatogenesis. Stem cell behaviour may have been fine-tuned to ensure the preservation of both fertility and genomic integrity over vastly differing life histories and reproductive timeframes and we hypothesise that the different transcriptomic and transitional states within the undifferentiated spermatogonial population provide a glimpse into the differing evolutionary strategies.

One interpretation of the data is that human SSCs have both ‘slow-cycling’ and ‘fast-cycling’ transcriptomic programs, shifting seamlessly from the former to the latter, a model not conceptually dissimilar to that of haemopoietic stem cells. Human HSCs comprise three subpopulations ^89^: dormant (long-term residency in G_0_), homeostatic/steady-state (occasionally entering the cell cycle and characterised by low metabolic activity), and activated (cycling continuously, and from which a proportion will differentiate). While it is in principle possible to project human HSCs onto the SPG atlas to determine whether dormant/homeostatic/active HSC subpopulations correspond to their equivalents in this system, in practice this is not a viable option. This is because HSC datasets are typically FACS-sorted on the basis of markers not widely expressed in undifferentiated spermatogonia, most notably *CD34* (which was only detected in < 5% of SSCs; **Supplementary Table 7**), leaving too few cells for a meaningful analysis. Nevertheless, it may still be instructive to draw a parallel between HSCs and human SSCs, in which state 0 comprise ‘homeostatic’ SSCs (and possibly also a subset transitioning to or from quiescence) and state 0A ‘activated’ SSCs (which have metabolically shifted to OXPHOS). While we concluded that the state 0 cluster was not as a whole quiescent, we could not rule out the possibility that some cells had a direct relationship to an undefined and hard-to-isolate quiescent state. While we also found no obvious analogue of the ‘dormant’ HSC state in SSCs, the transcriptional program of state 0B appears ‘suppressive’, which may be functionally equivalent.

Another stem cell system that may parallel the strategies adopted by SSCs is that of neural stem cells (NSCs). We have shown above that there is little transcriptomic evidence of undifferentiated spermatogonia showing G_0_-quiescence. However, while ‘quiescence’ is conventionally equated with G_0_ (by analogy to a state of nutrient withdrawal-induced arrest in yeast ^90^) this has been challenged by the finding that in *Drosophila*, quiescent neural stem cells arrest heterogeneously, with 75% residing in G_2_ instead ^91^. Similarly, mice have two types of quiescent NSC – ‘dormant’ (which have not previously proliferated) and ‘resting’ (which have proliferated at least once before), analogous to G_0_ and G_2_-quiescence, respectively – with the latter reactivating more quickly ^92^. An alternative interpretation of our analysis is that, like NSCs, SSCs also arrest heterogeneously. Consistent with this, a high proportion of cells in state 0 are in G_2_/M (48%; **Supplementary Table 13**) although also of note is that STK11/LKB1, which is implicated in establishing germline G_2_-quiescence in the ‘dauer’ state of *C. elegans* ^93,94^, is differentially expressed only in state 0 (**Supplementary Table 7**). Interestingly, state 0B appears more likely than state 0 to be held at G_2_ as it not only shows a higher proportion of cells in that phase (67%, the highest proportion of G_2_/M cells for all three undifferentiated spermatogonial states; **Supplementary Table 13**) but has a plausible biological reason to be arrested at that point, namely DNA repair to maintain germline integrity (high-fidelity double-strand break repair can only occur during the S/G_2_ phases because it requires a homologous repair template ^92^). Indeed, each of states 0, 0A, and 0B are skewed towards G_2_/M and away from G_1_, although state 0B most of all: only 5% of its cells are positioned in G_1_, in contrast to 9% for state 0A and 12% for state 0 (**Supplementary Table 13**; see also **Supplementary Text**). The relative proportion of G_2_/M to S is also seemingly more skewed for 0B than the other states. The G_2/_M to S ratio for states 0 and 0A approximates parity whereas for state 0B it appears skewed by 2:1 towards G_2_/M (**Supplementary Table 13**). A prolonged G_2_ phase has been associated with regeneration in other systems (including muscle stem cells in zebrafish embryos ^95,96^, adult stem cells in the intestinal crypts of the naked mole rat, the longest-living rodent ^97^, and adult stem cells of the regenerative *Hydra* polyp, which alternate between G_2_ and S, with minimal G_1_ ^98^), of note here given the presumably high selective pressure on SSCs to maintain a low germline mutation rate. A conceptual advance offered to SSC biology by NSCs is in suggesting that a reserve population (if present) may not be ‘dormant’ or ‘outside the cell cycle’ at all but ‘resting’, ‘waiting’ or ‘paused’; that is, not compromising the ability to be ready-to-react (parallels between NSCs and SSCs are discussed further in the **Supplementary Text**).

With the transcriptomic programs of undifferentiated spermatogonia broadly outlined, our most notable finding is that in a cross-species comparison, states analogous to human states 0 and 0B were absent from adult rats and mice, which suggests that the functions they implement are not required. We should recall at this point that not only are the expression profiles of states 0 and 0B slightly more correlated with each other than either are to state 0A (**Supplementary** Figure 10) but that among their set of differentially expressed genes, those GO terms which distinguish them have lower support (**Supplementary Table 11**). In this respect, and given states 0 and 0B are ‘not activated’ (i.e., not showing OXPHOS metabolism), they may have a particularly entwined set of functions. As a program of coordinated expression but unclear function, we conjectured state 0B is ‘suppressive’ both because it is (in part) transcriptionally directed towards state 0 (**Supplementary** Figure 11) and because during early development the function of its main diagnostic marker, *NANOS2*, is to induce mitotic arrest and degrade RNAs involved in the induction of meiosis ^73^, in essence slowing down cellular maturation. Consistent with this, candidate target genes for NANOS2 include *Taf7l*, *Dazl*, *Sycp3*, and *Prdm9* ^70^, the former differentially expressed immediately downstream of state 0B, in state 0A/1, the middle two differentially expressed in the ‘late diff SPG’, ‘leptotene’ and ‘zygotene’ clusters, and the latter, a key regulator of meiotic double-strand breaks ^99^, differentially expressed only in ‘leptotene’ (**Supplementary Table 7**).

Nevertheless, the full suite of NANOS2 targets and interacting partners remain elusive, and its precise function in adults a current gap in our knowledge. For instance, a previous study in mice found that NANOS2 requires a germ cell-specific RNA-binding protein, DND1, to associate with its target RNAs ^100^; curiously, we found no evidence of its orthologue in the adult human germline (**Supplementary Table 7**).

However, assuming state 0B is a dedicated ‘suppressive program’, why would adult rodents not require this? Although the basic elements of spermatogenesis are conserved between rodents and humans ^6^, the degree of transcriptomic similarity between undifferentiated primate and rodent spermatogonia is relatively low ^5^. We have noted above that in mice, undifferentiated spermatogonia expressing *Nanos2* are most likely to also express *Gfra1* ^64^ and that accordingly their closest human counterpart resembles state 0A, not 0B. These *Gfra1*+ *Nanos2*+ cells self-renew while producing a differentiation-primed *Ngn3*+ *Miwi2*+ *Rarγ*+ population that differentiates without self-renewal (reviewed in ^101^), cells which should in principle be analogous to the human ‘early diff SPG’ cluster if the species’ trajectories were held in common. However, we can see from our data this is not the case: in human spermatogonia, *PIWIL4* (*Miwi2*) is differentially expressed in state 0 (i.e. downstream to where it is expressed in mice), *RARG* (*Rarγ*) shows no evidence of differential expression in any cluster of the SPG atlas, and *NEUROG3* (*Ngn3*) is not expressed at all (**Supplementary Table 7**). Other differences between the primate and rodent germline are apparent even from the perspective of germline specification. For example, in humans, SOX17 is a critical regulator of PGC fate ^102^ although this factor plays no role in establishing the germline of mice ^103^. Mice also maintain a proportionately smaller stem cell pool than humans although compensate with a greater number of transit amplification divisions, ultimately producing around ten times more sperm per gram of testis parenchyma per day ^6^. What may underpin these differences between human and rodents is their relative age at sexual maturity and lifespan. Using estimates of the average age of male sexual maturity from the AnAge database ^104^, it is apparent that mice are able to produce large volumes of sperm comparatively rapidly, entering puberty practically at birth and accomplishing in 2 months what in humans takes around 14 years (**Supplementary Table 18**). As such, it is possible that relatively longer-lived species retain into adulthood the otherwise foetal state 0B, so as to reduce mutagenic load over the long term while ensuring the regular production of gametes across their longer reproductive lives. Consistent with this, the germline mutation rate of humans is an order of magnitude lower than mice (the ‘modeled rate per year’ from ref. ^1^ is 3.52×10^−9^ for mice and 4.18×10^−10^ for human). Overall, we suggest that shorter-lived species have adopted a different evolutionary strategy for producing sperm – one where undifferentiated adult spermatogonia do not employ the state 0B program and there is little selective benefit in ‘waiting’.

## Materials and Methods

### Integrated scRNA-seq atlases of the post-pubescent human testes and spermatogonial stem cells

To generate an integrated single-cell atlas of human spermatogonial stem cells, we first obtained 34 publicly-available post-pubescent (14-66 years) scRNA-seq testes samples, representing data from 29 individuals across 9 studies ^10,15,16,20,22–26^. Data sources and sample metadata are detailed in **Supplementary Tables 1 and 2**, respectively. To mitigate batch effects, we excluded from consideration samples which did not use an Illumina sequencer (e.g. ^105^); sequenced too few cells for use with a common informatic workflow, detailed below ^11,42,44^ (< 100 cells; these samples are typically from studies employing the whole-transcript but low-throughput Fluidigm C1/SmartSeq or SmartSeq2 methods ^106^); used FACS-sorted, enriched, or otherwise pre-selected cell populations rather than enzymatically-digested testes tissue ^24,43^ (both because the selection process may activate the cells and because the meaningful integration of datasets would not be possible, these samples having biased cell populations); and those where the donor had sex chromosome aneuploidy (e.g. ^41^), testosterone-suppression, non-obstructive azoospermia, or otherwise impaired fertility.

For each sample, cell x gene count matrices were obtained using the kallisto/bustools (KB) v0.26.3 ‘count’ workflow ^39^ with parameter ‘--filter bustools’ and, as a transcriptomic index, human genome GRCh38.p13, obtained from Ensembl v104 ^107^ (two indexes were built using this genome, one for expression quantification and one for velocity analysis, discussed below). As an initial QC step for each sample, KB filters cells on the basis of a knee plot ^108^, retaining only those above a sample-specific, empirically-derived, inflection point. For each sample, we also retained only those genes detected in at least 3 cells, and only those cells where the total number of genes was > 1000 and < 10,000, the total number of UMIs was > 2000 and < 50,000 and the proportion of mitochondrial genes detected was < 5% of the total. Each count matrix was then processed using Seurat v4.0.3 ^40^ with the normalisation procedure SCTransform ^109^, setting method to ‘glmGamPoi’ ^110^ and mitochondrial gene content, total gene count, total RNA count and both ‘S score’ and ‘G2M score’ as regression variables (using 67 cell cycle phase-associated genes from ^111^ and listed in **Supplementary Table 4**). Finally, to produce a testis atlas of 60,427 cells, all 34 samples were integrated using Seurat’s rPCA ‘anchor’ technique with 5,000 features. Seurat recommends rPCA for large datasets and requires the user define a ‘mutual neighbourhood’ in which to search for integration anchors. For this purpose, we used as our ‘neighbourhood’ the combined set of 8 samples from the Di Persio ^15^, Sohni ^16^, and Zhao ^54^ studies, as together these contributed the greatest number of cells (**Supplementary Table 3**); 30,776 cells, or 51% of the total.

To interpret the structure of the testis atlas, transformed gene counts were used for principal component (PC) analysis, with elbow plots (which rank PCs by the percentage of variance explained) used to determine the optimal number of PCs for subsequent analysis, including – for visualisation – performing non-linear dimensional reduction using the UMAP method ^112^. The ‘elbow’ was programmatically derived on the basis of being either where the percent change in variation between consecutive PCs was less than 0.1%, or where the PC contributes < 5% of the variation and all previous PCs have cumulatively contributed > 90%, whichever is lower. We retained for analysis all PCs up to and including this elbow PC.

A k-nearest neighbour graph was then constructed using these top PCs, with community detection (i.e. clustering) performed using the Smart Local Moving algorithm ^113^ with resolution 0.15. This resolution was empirically chosen on the basis of both a Clustree v0.5.0 ^46^ dendrogram and manual review (using the interactive browser cellxgene v0.18.0 (https://github.com/chanzuckerberg/cellxgene)), with clusters made at resolutions in the range 0 to 0.5, at intervals of 0.05, interpreted in the context of known testes cell markers (given in **Supplementary Table 4** and sourced from ^10,16,25,41–44^) and, to facilitate an unbiased annotation of cluster identities, an all-against-all differential expression analysis.

To that end, we determined the top candidate marker genes per cluster using Seurat’s FindAllMarkers function with parameters min.pct = 0.25, logfc.threshold = 0.25 and only.pos = TRUE which, respectively, require that a gene be expressed in >25% of the cells in a given cluster, have a log_2_ fold-change difference in expression relative to all other clusters of > 0.25 (i.e. have 2-fold greater average expression) and to be a positive cluster marker, i.e. to be on average more highly expressed in that cluster than all others.

Finally, two clusters of undifferentiated SSC and differentiating spermatogonia (indicated on **Figure 1** as ‘undiff SPG’ and ‘diff SPG/early meiosis’, respectively) were subset and a higher-resolution ‘SPG atlas’ created by re-running the above QC, SCTransform and clustering steps on the subset cells. The data were initially clustered at very high resolution (5.0) such that cluster IDs could be assigned to three small contaminating cell populations; these were removed and the dataset iteratively re-integrated to converge on the final atlas. This process is further detailed within the scripts used for this analysis, available at github.com/sbush/spg_atlas. Note that because meaningful re-integration could not be made for samples with < 100 cells and that for any given testis sample, only a small proportion of its cells would be SSCs, the SPG atlas was restricted to data from 9 of the original 34 samples, representing 5 of the 9 studies (data from ^10,22,25,26^ was excluded).

### scRNA-seq analysis of non-adult human and non-human samples

Data from 7 foetal and 21 post-natal pre– and peripubertal human scRNA-seq samples (foetal age range 8-23 weeks; post-natal age range 2 days to 13 years) ^10,16,23,25,27–29^, and from 1 pig ^36^, 1 sheep ^37^, 2 buffalo ^38^, 4 cynomologus macaque ^30^, 11 rhesus macaque ^5,20^, 8 rat ^35^, and 37 mouse samples ^31–34^, were individually processed using the same KB/Seurat workflow, with the same parameters, described above. Data sources and sample metadata are detailed in **Supplementary Tables 1 and 2**, respectively.

### RNA velocity analysis

RNA velocity was calculated using scVelo v0.2.5 ^48^ after having first processed each sample using the KB v0.26.3 ‘count’ workflow ^39^ with parameters ‘––workflow lamanno ––filter bustools ––loom’. For each of the 9 samples in the SPG atlas, we then parsed the appropriate ‘counts_unfiltered/adata.loom’ file to extract its list of cell barcodes. We calculated velocity only for cells included in the final SPG atlas, projecting this onto the pre-existing UMAP embedding using the ‘velocity_embedding_stream’ command.

### Enrichment and impact analyses

For each cluster in the SPG atlas, GO term and KEGG pathway enrichment was assessed using the R/Bioconductor packages topGO v2.22.0 ^114^ and SPIA (Signalling Pathway Impact Analysis) v2.46.0 ^115^, respectively. The topGO package employs the ‘weight’ algorithm to account for the nested structure of the GO tree with correction for multiple hypothesis testing intrinsic to the approach ^114^. This requires a reference set of GO terms, built manually from GRCh38.p13/Ensembl v104 and filtered to remove those terms with evidence codes NAS (non-traceable author statement) or ND (no biological data available), and those assigned to fewer than 10 genes in total.

Significantly enriched GO terms (p < 0.05) were reported only if the observed number exceeded the expected by 2-fold or greater.

To run SPIA, we used the set of human KEGG pathways ^116^ as of 8^th^ March 2022 (available at https://www.genome.jp/kegg-bin/download_htext?htext=br08901.keg). SPIA combines evidence from both a classical enrichment analysis and a bootstrap procedure to quantify the differential impact upon a given pathway. The former considers each pathway a gene list and gives no weight to topology, that is, to the relative position of each gene within that pathway ^117^. The latter measures the perturbation on a given pathway by propagating fold changes in expression across its topology. A global probability value, P_G_, is then calculated for each pathway incorporating both the log fold-change of the differentially expressed genes, the statistical significance of the enrichment of any pathway and the topology of the pathway itself. Consequently, SPIA assess not only which pathways are enriched for DE genes, but whether the DE is likely to impact upon the function of that pathway. For example, if those DE genes assigned to a pathway are among the downstream genes of that pathway, changes in their expression levels are less likely to affect the pathway as a whole. We retain only those pathways with P_G_ < 0.05 after controlling for a FDR of 5%.

### Projection of individual samples onto the SPG atlas

Individual samples, from both human and non-human samples, were projected onto the human whole-testis and SPG atlases using the ‘FindTransferAnchors’ and ‘MapQuery’ commands of Seurat v4.0.3 ^40^, and following a two-step strategy described below. To facilitate projections, gene names were standardised using one-to-one orthology relationships obtained from Ensembl BioMart ^107^ release 104 (genome versions GRCm39, mRatBN7.2, GCA_003121395.1, Oar_rambouillet_v1.0, Macaca_fascicularis_6.0, Mmul_10, and Sscrofa11.1, for mouse, rat, buffalo, sheep, cynomolgus macaque, rhesus macaque, and pig, respectively). Cells from a given sample were then projected onto the whole-testis UMAP so as to distinguish germline from somatic cells. Only these putative germ cells were then projected onto the SPG atlas.

Seurat’s projection algorithm works by assigning a mapping score to each cell, from 0 to 1 (lowest to highest), first by projecting the query cell into the reference space, then projecting that cell back onto the query, and then determining to what extent the local neighbourhood was altered in the process (detailed further at https://rdrr.io/cran/Seurat/man/MappingScore.html). For human and mouse samples, we required that each cell was projected with a minimum score of 0.8, thereby restricting analysis to cells with high transcriptomic similarity (cells not meeting this threshold would not be shown in the projection figures). For the other species (sheep, pig, buffalo, both macaques, and rat), whose genomes are generally not as well annotated as these models, we varied projection score from 0 to 0.8 at intervals of 0.2. A projection score of 0 ensured each cell would be plotted according to its ‘best match’ on the SPG atlas, even if in absolute terms that match was poor.

## Supplementary Figure Legends

**Supplementary Figure 1.** No overt batch effects in the adult testis atlas on the basis of study of origin (A), donor age (B), or sample accession (C).

**Supplementary Figure 2.** No overtly disproportionate contribution made by individual samples to the adult testis atlas, on the basis of number of genes per cell (A), number of UMIs per cell (B), and percentage of mitochondrial genes per cell (C).

Boxes represent the interquartile range of each variable, with midlines representing the median. Upper and lower whiskers extend, respectively, to the largest and smallest values no further than 1.5× the interquartile range. Data beyond the ends of each whisker are outliers and plotted individually.

**Supplementary Figure 3.** No overt batch effects in the adult SPG atlas on the basis of study of origin (A), donor age (B), or sample accession (C).

**Supplementary Figure 4.** No overtly disproportionate contribution made by individual samples to the adult SPG atlas, on the basis of number of genes per cell (A), number of UMIs per cell (B), and percentage of mitochondrial genes per cell (C).

Boxes represent the interquartile range of each variable, with midlines representing the median. Upper and lower whiskers extend, respectively, to the largest and smallest values no further than 1.5× the interquartile range. Data beyond the ends of each whisker are outliers and plotted individually.

**Supplementary Figure 5.** Proportional contribution of UMIs made by each (A) sample and (B) study to each cluster in the adult SPG atlas.

**Supplementary Figure 6.** Reciprocal projection of (A) the ‘undiff SPG’ and ‘diff SPG’ clusters of the whole-testis atlas onto the SPG atlas, and (B) the SPG atlas onto the whole-testis atlas, to confirm that the two atlases shared the same cell populations.

**Supplementary Figure 7.** Relative expression, across the SPG atlas, of key genes spanning the temporal order of gametogenesis.

This figure shows the relative expression, across the SPG atlas, of 16 genes using a yellow (minimum) to red (maximum) colour gradient. Note that each plot has been individually scaled to the maximum expression value of that gene and so the colour scales are not comparable across plots (and so are not shown). For illustrative purposes, it suffices to note that the maximum expression of each gene is confined to a particular cell type, and is widely considered a discriminative biomarker of it (as detailed in **Supplementary Table 4**).

**Supplementary Figure 8.** Clustree ^46^ plot showing the effect of different resolutions on the cluster composition of the SPG atlas. Clustering was performed using Seurat with resolution varied from 0 to 2 at 0.1 intervals. Seurat labels clusters according to their size, with “cluster 0” the largest. The set of nodes on each row represent the clusters formed at each resolution (i.e. at resolution 0, there is one node, and at resolution 0.1, three nodes). Arrows between nodes indicate their relationship to clusters formed at the next level of resolution. For example, as resolution increases from 0.2 to 0.3, the “cluster 0” node is subset into two nodes, “cluster 1” and “cluster 2”, indicating that the cluster has been broken up. The relative size, colour and thickness of each arrow indicate the proportion of cells contributed to each cluster.

**Supplementary Figure 9.** Single-cell expression atlas of spermatogonia and their progenies, at Seurat clustering resolutions (A) 0.3, (B) 1.1, and (C) 1.2, these partitioning the undifferentiated spermatogonial compartment into two, three, and four subsets, respectively. Note that panel B is identical to **Figure 1B**.

**Supplementary Figure 10.** Relative expression of genes in each of the three undifferentiated spermatogonial clusters (states 0, 0A/1, and 0B), using the SPG atlas clustered at resolution 1.1. The figure plots, per gene, the mean expression across all cells in the state 0 cluster relative to all cells in the state 0A/1 cluster (red circles), and to all cells in the state 0B cluster (blue triangles), demonstrating a substantial overlap between the three. There are significantly high Spearman’s correlations between all three comparisons (*rho* for state 0 vs. state 0A/1 = 0.934; for state 0 vs. state 0B = 0.955; for state 0A/1 vs. state 0B = 0.939; p < 2.2×10^−16^ in each case). Expression levels are quantified by Seurat and have arbitrary units. Raw data for this figure is available in **Supplementary Table 7**.

**Supplementary Figure 11.** Visualisation of the results of an scVelo RNA velocity analysis on the SPG atlas UMAP.

The streamlines indicate developmental trajectories, with the direction and thickness of each representing the orientation of the trajectory and the magnitude of its velocity, respectively. Strong and unidirectional streamlines are most notable in the ‘early diff SPG’ cluster, proceeding leftwards to the onset of meiosis. There are similarly strong directional streamlines proceeding through the early stages of prophase (where the trajectory is terminated) although nevertheless a number of backward-pointing arrows between stages, interpreted here to indicate the activity of the meiotic checkpoint network ^44^ that regulates progression. Moving to the right of the figure, we can see that the directionality of the cells is less certain. Between the undifferentiated spermatogonial ring and the ‘early diff SPG’ cluster, there are multiple clear streamlines pointing both up (towards differentiation) and down, consistent with previous suggestions that prior to differentiation, undifferentiated spermatogonia are not irreversibly committed to their fate and may instead show more plastic behaviour ^10^. Streamlines within the ring are discussed in the main text.

**Supplementary Figure 12.** Verification of the two-step projection strategy. (A) Projection of three FACS-sorted steady-state spermatogonial samples (from ^24^) onto the adult testis atlas, so as to distinguish germline from somatic cells. (B) Projection of these putative germ cells onto the SPG atlas.

**Supplementary Figure 13.** Projection of rhesus macaque germ cells onto the SPG atlas, part 1 of 2. Details of the samples included in this analysis are given in **Supplementary Table 2**. The number of cells projected onto each cluster, for a range of minimum projection scores (0 to 0.8 at 0.2 intervals), are given in **Supplementary Table 16**, with this figure plotted with no minimum threshold required.

**Supplementary Figure 14.** Projection of rhesus macaque germ cells onto the SPG atlas, part 2 of 2. Details of the samples included in this analysis are given in **Supplementary Table 2**. The number of cells projected onto each cluster, for a range of minimum projection scores (0 to 0.8 at 0.2 intervals), are given in **Supplementary Table 16**, with this figure plotted with no minimum threshold required.

**Supplementary Figure 15.** Confirmation that human states 0 and 0B-like populations are not obviously apparent in adult mice. (A) Projection of two sorted populations of undifferentiated mouse germ cells (from ^85^) onto the adult testis atlas, so as to distinguish germline from somatic cells. (B) Projection of these germ cells onto the SPG atlas. These two samples are biological replicates from 8-10 week old mice and represent cells isolated from the testes on the basis of *Plzf* (ZBTB16) gene expression, and then stained for CD9 and c-Kit, markers of stem and differentiating spermatogonia, respectively. As both CD9 and ZBTB16 are detected in a comparably high number of cells in each of human states 0, 0A and 0B (**Supplementary Table 7**; note that ZBTB16 is differentially expressed in state 0, too), we considered it reasonable to perform a mouse-to-human projection. Note that CD9 is also expressed by somatic cells and so a small proportion of each sample are excluded by the two-step projection strategy (because some cells project to the myoid/Leydig cluster rather than the germline).

**Supplementary Figure 16.** Presence of human states 0 and 0B-like populations in embryonic mice.

(A) Projection of three populations of embryonic day 16.5 mouse PGCs (from ^86^) onto the adult testis atlas, so as to distinguish germline from somatic cells. (B) Projection of these germ cells onto the SPG atlas.

## Supplementary Tables

**Supplementary Table 1.** Source of testes scRNA-seq datasets.

**Supplementary Table 2.** scRNA-seq sample metadata.

**Supplementary Table 3.** Number of cells, genes, and average genes per cell, per sample.

**Supplementary Table 4.** Cell and spermatogonial stem cell state biomarkers.

**Supplementary Table 5.** Adult human testis expression atlas.

**Supplementary Table 6.** Adult human SPG expression atlas (Seurat clustering resolution 0.3).

**Supplementary Table 7.** Adult human SPG expression atlas (Seurat clustering resolution 1.1).

**Supplementary Table 8.** Adult human SPG expression atlas (Seurat clustering resolution 1.2).

**Supplementary Table 9.** Enriched biological process GO terms for genes differentially expressed in the adult testis atlas.

**Supplementary Table 10.** Enriched biological process GO terms for genes differentially expressed in the SPG atlas (in an all-against-all analysis).

**Supplementary Table 11.** Enriched biological process GO terms for genes differentially expressed in the SPG atlas (in an X-against-Y analysis).

**Supplementary Table 12.** Enriched KEGG pathways for genes differentially expressed in the SPG atlas (in an X-against-Y analysis).

**Supplementary Table 13.** Proportion of cells in each SPG atlas cluster in each cell cycle phase.

**Supplementary Table 14.** Expression of quiescence-associated genes in undifferentiated spermatogonial states 0, 0A and 0B.

**Supplementary Table 15.** Proportion of cells from 62 human samples projecting to each of seven SPG atlas clusters.

**Supplementary Table 16.** Proportion of cells from 1 sheep, 1 pig, 2 buffalo, 4 cynomolgus macaque, 11 rhesus macaque, and 8 rat samples projecting to each of seven SPG atlas clusters.

**Supplementary Table 17.** Proportion of cells from 37 mouse samples projecting to each of seven SPG atlas clusters.

**Supplementary Table 18.** Average age of sexual maturity and maximum lifespan.

**Supplementary Table 19.** Proportion of cluster-enriched genes, from Huang, *et al.* (2023), detectable in the SPG atlas.

**Supplementary Table 20.** Proportion of core genes, from Murat, *et al.* (2023), detectable in the SPG atlas.

**Supplementary Table 21.** Gene expression in mouse germ cells from postnatal days 5 and 6.

**Supplementary Table 22.** Cell cycle phase predictions for the cells comprising the SPG atlas.

## Institutional Review Board Statement

Ethical review and approval were not applicable for this study.

## Informed Consent Statement

Not applicable for this study.

## Acknowledgements

We would like to thank Laurence Brown and the University of Oxford Interactive Data Network for support in hosting the SPG atlas ShinyApp.

## Data Availability

All raw sequencing data was sourced from openly available databases using the accession numbers given in **Supplementary Table 1**. An interactive version of the SPG atlas, made using ShinyCell ^118^, is available at https://trainingidn.shinyapps.io/hu_spermatogonial_atlas_shinyApp.

## Code Availability

All scripts used to perform the analyses and create both figures and tables are available at www.github.com/sjbush/spg_atlas.

