## Supplementary Text for "Human spermatogonial stem cells retain states with a foetal-like signature"

This supplementary text elaborates on several findings described in the main text, namely:

- Orthogonal validation of the SPG atlas clusters
- Characteristics of state 0B other than *NANOS2* upregulation
- Seemingly fewer human-mouse orthologues in state 0 than in states 0A or 0B
- The first mouse germ cell population to emerge after birth appears PGC-like
- Re-assessing cell cycle phase using different gene marker sets
- The human state 0 transcriptomic program resembles, in part, that of mouse neuroblasts
- Projection of a time course of mouse germ cells onto the SPG atlas
- Projection of a time course of human germ cells onto the SPG atlas

***Orthogonal validation of the SPG atlas clusters***

A recent multi-omics study combined scCOOL-seq and scRNA-seq (specifically, STRT-seq) to simultaneously capture the transcriptome, methylome and chromatin accessibility state of the same adult human testicular cell ^1^. Clustering of their STRT-seq data produced a transcriptomic atlas of 14 germline and 3 somatic cell types, capturing an average of 9323 genes per cell for 1097 cells (approx. three times as many genes/cell than in our own atlas, but 60-fold fewer cells; **Supplementary Table 3**). Our seven scRNA-seq clusters (at resolution 1.1) were ‘state 0’, ‘state 0A/1’, ‘state 0B’, ‘early diff SPG’, ‘late diff SPG’, ‘leptotene’ and ‘zygotene’ (**Supplementary Table 7**), whereas the corresponding Huang, *et al.* STRT-seq clusters were labelled ‘undiff.SPG-1’, ‘undiff.SPG-2’, ‘diff.ing SPG’, ‘pre-leptotene’, ‘leptotene-1/2’, ‘leptotene-3’, and ‘zygotene’, on the basis of their names alone showing comparatively greater resolution for primary spermatocytes but lower resolution for undifferentiated, differentiating, and differentiated spermatogonia. Although using a different scRNA-seq technology to those in the present study, the cluster markers for undiff.SPG-1 (*PIWIL4*+ *ID4*+ *L1TD1*-) show good correspondence to those of both states 0 and 0B, undiff.SPG-2 (*PIWIL4*- *L1TD*+) to state 0A/1, and diff.SPG (*KIT*+ *MKI67*+) to early diff SPG: *ID4* is differentially expressed in our states 0, 0A/1, and 0B, *PIWIL4* in both states 0 and 0B, and *L1TD1* and *MKI67* only in state 0A/1 and early diff SPG, respectively (**Supplementary Table 7**).

Huang, *et al.* refined their SSC/SPG clusters by incorporating chromatin accessibility data, identifying 3353 distal *cis*-regulatory elements corresponding to 2529 genes. Clustering analysis of this data produced five clusters, A1 through A5, which share identities with their undiff.SPG-1 (A1), undiff.SPG-2 (A3) and the diff.ing SPG (A5) clusters, as well as with their intersections: A2 sharing similarity with both undiff.SPG-1 and undiff.SPG-2, and A4 with both undiff.SPG-2 and diff.ing SPG. As there are differences in stochastic capture by both gene expression and chromatin accessibility datasets – the majority of genes were not found in common – the latter also serves as an independent validation set to assess our cluster identities. Accordingly, we determined the proportion of genes in each of clusters A1 to A5 that were also differentially expressed in our seven clusters. We found that the highest overlaps were of A1 (*CELF4*+ *KLF6*+ *POU3F1*+ *TCF3*+) and A2 (*BMPR1A*+ *FGFR3*+ *ZBTB16*+) with state 0 (with the exception of *BMPR1A*, all seven genes are DE in state 0), of A3 (*RET*+) with state 0A/1, and of both A4 and A5 (*NANOS3*+ *DMRTB1*+) with early diff SPG (in which both genes are differentially expressed; **Supplementary Table 19**).

On the basis of these results, one-to-one correspondences of ‘state 0’ with both ‘undiff.SPG-1’ and ‘A1’, and ‘state 0A/1’ with both ‘undiff.SPG-2’ and ‘A3’ appear evident. Although no clear correspondence could be found with our state 0B cluster, by considering the differential expression of its marker genes, it appears subsumed into undiff.SPG-1 and therefore more similar to state 0 than to state 0A/1, consistent with our own results (discussed in the main text in relation to **Supplementary Figure 10** and **Supplementary Table 11**). Accepting a correspondence between the Huang clusters and our own, we could use the findings of their study to refine our functional interpretations of state 0. To that end, supporting our inference of relative hypoxia in state 0, GO terms enriched among the ‘A1’ set of open chromatins include both “regulation of glycolytic process” and its parent term, “regulation of generation of precursor metabolites and energy.” More tenuously, if assuming undiff.SPG-1 overlaps both state 0B and ‘A2’, then consistent with the ‘suppressive’ role of the former (discussed further below) is a GO term enriched in A2: “negative regulation of mitotic cell cycle phase transition.”

***Characteristics of state 0B other than NANOS2 upregulation***

State 0B, while typified by, is not simply synonymous with higher *NANOS2* expression. For instance, if ranking the 277 genes differentially expressed (DE) in state 0B by their average log_2_ fold change, there are six genes more highly upregulated than *NANOS2*: in order, *CCDC106*, *RPL3*, *SCAMP2*, *XAB2*, *SGF29*, and *TMEM39B* (**Supplementary Table 7**). If ranked instead by the absolute difference in the proportion of cells expressing this gene in state 0B, relative to the proportion of cells expressing this gene in all other clusters, then 19 genes show higher disparities than *NANOS2*, the top four of which are, in order, *CCDC106*, *TMEM39B*, *SCAMP2*, and *SGF29*. Arguably, these genes contribute as substantively to the ‘state 0B’ expression profile as *NANOS2*, although this distinguishes itself by being – unlike either of the above – almost exclusively expressed in state 0B instead; it is detectably expressed in only 2.7% of the other cells in the SPG atlas (**Supplementary Table 7**).^[1]*^

On the basis of both the functions associated with selected DE genes, and the projection of mitotically-arrested foetal germ cells exclusively onto the ‘state 0B’ cluster (**Figure 2A** and **Supplementary Table 15**; see also below), we conjectured that state 0B was a transcriptomic program for actively suppressing proliferation. For this hypothesis to hold, one would expect many genes characteristic of state 0B to function accordingly, with roles in cellular homeostasis or cell cycle progression, if not explicitly in mitotic arrest.

Critical to cellular homeostasis and the maintenance of genomic integrity is the tumour suppressor p53 ^2^, which accumulates in response to, among others, DNA damage and hypoxia (in relative conditions of which we presume state 0 cells, which are transcriptionally ‘adjacent’ to 0B, reside). Active p53 arrests the cell cycle to ensure potentially damaged cells do not multiply (it does so by upregulating expression of *CDKN1A*/p21, facilitating the formation of a protein complex which binds the promoters of many cell cycle genes, downregulating their activity; see review ^3^). Strikingly, the most highly upregulated gene in state 0B, *CCDC106,* has been shown in yeast to promote the degradation of p53 ^4^, with overexpression in mice associated with tumour cell proliferation ^5^. We cannot easily reconcile these observations with the hypothesis that state 0B is generally ‘suppressive’, other than by suggesting that the nature and extent of its cell cycle arrest could be very precisely regulated, therefore not only positively but negatively (i.e. by *CCDC106*). Interestingly, however, the second most highly upregulated gene in state 0B, *RPL3* (already discussed in the main text in the context of its involvement in DNA repair ^6^) complements *CCDC106* in function: it can induce p21-dependent cell cycle arrest and apoptosis in the absence of p53 ^7^.

Other genes upregulated in state 0B are more obviously consistent with a ‘suppressive’ hypothesis. For instance, *SGF29* is functionally involved in cellular senescence in human mesenchymal progenitor cells, with its overexpression associated with reduced cellular proliferation rates, elevated p21 levels, and an increased DNA damage response ^8^. The multi-functional protein *XAB2* also has a role in mitotic cell cycle regulation, such that knockdown results in mitotic arrest (at G_2_/M phase) and catastrophe ^9^ (delayed mitotic-linked cell death resulting from premature or inappropriate entry into mitosis ^10^). Also with a key role in cellular homeostasis, *SCAMP2* (a conserved membrane protein which acts as a carrier in post-Golgi recycling pathways) has a prognostic association with acute myeloid leukemia, a malignant clonal disease of haematopoietic tissue characterised by dysregulated cell proliferation and apoptosis ^11,12^.

We argued in the main text that state 0B had its strongest associations with cell cycle regulation and DNA repair, which we can also support by a more general interpretation of its GO terms. As context, it is immediately apparent that all three undifferentiated spermatogonial clusters (states 0, 0A/1, and 0B) are replete in ‘regulation’ related terms, both positive and negative, suggesting the precise coordination of each state (**Supplementary Tables 10 and 11**). Of particular note, however, is “negative regulation of ubiquitin protein ligase activity” in state 0B, a relatively specific GO term (being assigned only to 13 genes) but one where the number of observed DE genes far exceeds the expected (54% of the total genes with this term are DE in state 0B) (**Supplementary Table 10**). Ubiquitin, a post-translational modification that tags proteins for degradation, is central to every process in which the speculated function of state 0B resides: to DNA repair, the maintenance of genomic integrity, cell cycle checkpoints, and programmed cell death ^13^. Related to this, in the pairwise comparison of state 0B with state 0 (**Supplementary Table 11**), both “protein neddylation” and “protein deneddylation” appear as enriched GO terms (referring to the activity of NEDD8, a ubiquitin-like protein), both of which are also assigned only to a small number of genes (**Supplementary Table 11**). Neddylation dynamics play a critical role in the DNA damage response, in particular double-strand break repair (reviewed in ^14,15^), with deneddylation regulating the choice of repair pathway, decreasing its use of non-homologous end joining (NHEJ) in favour of homologous recombination ^16^. Interestingly, this observation could be related to low germline mutation rates: NHEJ, a comparatively ‘fast and simple’ repair process, is relatively imprecise (it has no proofreading activity) and so contributes to mutations that arise over time ^17^.

We have also noted that mitotically-arrested foetal germ cells project exclusively onto the ‘state 0B’ cluster (**Figure 2**) and that therefore they may implement similar transcriptomic programs. Using an integrated single-cell dataset comprising nine male embryos/foetuses (from 6 to 23 weeks post-conception), ^18^ identified a cluster of mitotically arrested foetal germ cells, characterised by the differential expression – relative to several other clusters of foetal germ cells – of 28 genes. Supporting a link between the ‘arrest’ and ‘0B’ programs, we found that 19 of these genes (68%) were differentially expressed in state 0B (*BBX*, *DCAF4L1, EGFL7, FHL1, HDAC5, HECTD1, ID1, ID4, KLHL35, PABPC4, PLD3, PODXL2, POLR2A, PTOV1, RRBP1, TNRC6B, USP11, VAMP2, ZBTB43*; **Supplementary Table 7**) although note that none were only differentially expressed in this state; rather, 16 of the 28 (57%) were also differentially expressed in state 0A and 15 (54%) in state 0. Collectively, these associations – while hardly definitive evidence that state 0B is ‘suppressive’ – suggest the hypothesis may at least merit consideration.

***Seemingly fewer human-mouse orthologues in state 0 than in states 0A or 0B***

One of our principal findings is that states 0 and 0B are present in adult humans, but not rodents. While our interpretation of this centred on the role key genes played in each state (notably *NANOS2*), a simpler hypothesis is that these states are absent from rodents because they disproportionately express human-specific genes. Consistent with this, of the 362 genes only DE in state 0, 81 (22.4%) had no orthologue in mouse, although the majority of these (62) were lncRNAs (**Supplementary Table 7**). This appears in contrast to states 0A/1 and (unexpectedly) 0B, the genes in which in principle both shared greater homology with mouse. Of the 324 genes only DE in state 0A/1, 27 (8.3%) had no orthologue in mouse, and of the 277 genes only DE in state 0B, 21 (7.6%) had no orthologue in mouse. In the latter case, 12 of these genes were lncRNAs, 3 were transcribed pseudogenes, and only 6 protein-coding: *BCAS4*, *FIP1L1*, *IFI27L1*, *PAGE2*, *RPL9*, and *SETD1A*^[2]*^ (although nevertheless *FIP1L1*, *PAGE2* and *RPL9* were each expressed in > 80% of the cells in the 0B cluster) (**Supplementary Table 7**).

Moreover, a previous study ^20^ identified 416 genes with conserved germline expression across 9 mammals (human, chimpanzee, bonobo, gorilla, gibbon, macaque, marmoset, mouse, and opossum), this set likely representing ancestral members of the core mammalian spermatogenic program. As expected, virtually all (>99%) of these ‘core’ genes were detectably expressed in every cluster of the SPG atlas (**Supplementary Table 20**), although we also noted that more were differentially expressed in state 0A/1 (n = 33) than in states 0 (n = 16) or 0B (n = 21). Although not formally assessed, it is possible that this represents a comparative relaxation of constraint on genes more strongly expressed in states 0 and 0B than 0A. This could contribute to the increased divergence of these states between species, consistent with the rapid evolution of the testis at the molecular level, ostensibly due to evolutionary pressure to be reproductively successful ^21^. That proportionately more of the ‘core’ genes appear differentially expressed in state 0A (and few genes differentially expressed in this state have no orthologue in mouse) is consistent with its ‘active’ – and presumably more highly conserved – role, of being inclined towards OXPHOS metabolism and commitment to differentiation.

***The first mouse germ cell population to emerge after birth appears PGC-like***

In mice, progressive re-entry of primordial germ cells (PGCs; also known as prospermatogonia or gonocytes) into the cell cycle commences around postnatal (P) days 1 to 3, with this process transitioning cell fate from PGC to SSC and establishing a foundational SSC pool for eventual steady-state spermatogenesis. In rodents a unique first wave of spermatogenesis occurs immediately after cell cycle re-entry and which emanates from a subset of non-self-renewing spermatogonia ^22^. This first wave differs from the steady-state of adulthood, which by definition requires a sustainable balance between SSC self-renewal and the initiation of differentiation.

This first wave is tightly synchronised, with key time points for the sequential appearance of particular cell types well defined. For instance, at P6 and P7, the seminiferous tubules only contain Sertoli cells and spermatogonia; accordingly, the P6 sample projected onto the SPG atlas in **Figure 4** only contains spermatogonia. Early spermatocytes (leptotene) first appear c. P9 and the pachytene stage of the first meiotic prophase initiates c. P14 ^23^ (with some minor variability in timing by species or strain of rodent ^24^), with the **Figure 4** projections at P10 and P15 also consistent with this.

We have also shown in **Figure 4** that the first mouse germ cell population to emerge after birth (detectable at P5), and which presumably initiates this first wave, does not resemble any of the human undifferentiated spermatogonial clusters; rather, it projects almost entirely to the human ‘early diff SPG’ cluster. This population is of particular interest as it is generally unclear as to what extent the transcriptomic profiles of first-wave germ cells differ from those which sustain steady-state spermatogenesis, and how the latter is established ^25^. It has previously been suggested that the foundational (steady-state) mouse SSCs maintained until adulthood resemble the ID4+ spermatogonia detectable at P6 ^25^, which is consistent with our projection of an independent sample at that time point (**Figure 4**), in which each of the human-like ID4+ undifferentiated spermatogonial clusters (states 0, 0A and 0B) could be detected.

As P5, but not P6, germ cells primarily resemble human ‘early diff SPG’, one interpretation of this is that a subset of this population are functionally committed and will take further developmental steps in the direction of meiosis (to produce the first wave of sperm), but that another subset must later self-renew, establishing the foundational SSC pool for steady-state spermatogenesis (which becomes visible from P6 onwards). This is consistent with the argument that the first wave of spermatogenic cells are not necessarily derived from SSCs (reviewed in ^22^) but from PGCs directly, implementing a distinct program that lacks the self-renewal stage ^26^. That this first germ cell population to emerge after birth serves a dual role would explain observations of the relative inefficiency of the first wave ^27^.

Accordingly, a fuller characterisation of this P5 population may give insight into which genes underpin the cell-fate transition from mouse PGC to SSC, and which ultimately establish the foundational population of SSCs for steady-state spermatogenesis. To explore this further, we return to the set of mouse cells projected to the SPG atlas in each of three samples shown in **Figure 4**, at P5 (ERS3000379, n = 282 cells; and ERS3000380, n = 474 cells) and P6 (SRS3990943, n = 820 cells). Note that to maximise the number of cells in each sample, each of these projections was made at a minimum confidence threshold of 0 (**Supplementary Table 17**). We then determined what proportion of cells in each sample contained detectable expression (≥ 1 read) of any given gene. As expected, the correlation between these two sets of values (that is, the transcriptomic profiles at each day) were near-identical for the two P5 replicates (Spearman’s *rho* = 0.99, p < 2.2e^-16^), but differed between the mean of the P5 replicates and the value for P6 (Spearman’s *rho* = 0.95, p < 2.2e^-16^). We then determined the difference between the proportion of cells in which a gene was expressed at P5 compared to P6, and vice versa, using this as a simple means of prioritising candidate genes that may distinguish the two. Although this approach can offer only a crude overview – no formal statistical approach was undertaken, and the sample size is limited – it may nevertheless highlight possible directions for further enquiry. It was first apparent that both the P5 and P6 populations had approximately comparable levels of the prototypical undifferentiated mouse spermatogonial markers *Gfra1* (c. 20% of cells in both the P5 and P6 samples), *Id4* (c. 40%), *Piwil4* (c. 10%) and *Rhox10* (c. 25%). Another characteristic marker of undifferentiated (but not differentiating) mouse spermatogonia is *Neurog3* ^28^. A previous study found that approximately 60% of sperm produced during the first wave of spermatogenesis originated from a *Neurog3*- population of germ cells, and that consequently these sperm must have bypassed the ‘undifferentiated spermatogonial’ stage and arisen directly from PGCs ^26^. Consistent with this, we found that of the two P5 samples, one had no detectable expression of *Neurog3* and the other found *Neurog3* in only < 2% of its cells. By contrast, at P6, *Neurog3* was detected in approximately 5% of cells.

Possible lines of enquiry may centre on those genes with greatest difference in proportional detection between P5 and P6 (**Supplementary Table 21**). Strikingly, a gene detected in c. 66% of P5 germ cells but only c. 12% of P6 germ cells was the classical PGC marker *Nanog* ^29^, implicating the P5 population as being potentially more ‘PGC-like’ than that of P6. A previous study of the neonatal (2-7 day old) human testis also identified a ‘PGC-like’ (*NANOG*+ *POU5F1*+) subset of germ cells, posited as a transitional stage between embryonic PGCs and adult SPG ^30^.

As we have argued in the main text, one of the principal differences between human and rodent spermatogonia is that the transcriptional programs implemented by the latter do not result in germ cells ‘waiting’ (consistent with which, the overall duration of spermatogenesis – that is, of differentiation from spermatogonia to mature spermatozoan – is c. 35 days in mouse ^31^ compared to 74 days in human ^32^). This point of transition from PGC to SSC would presumably determine how these foundational transcriptional programs for steady-state spermatogenesis are established, and in what way the two species differ. As such, it could be of interest to further interrogate the overlap between human PGC-like marker genes ^30^ and those comparatively enriched at mouse P5. Of immediate note is that a number of genes more highly differentially expressed in the human ‘PGC-like’ cluster (from ^30^) are also more highly detected in P5 germ cells relative to P6, reinforcing the annotation of a conserved ‘PGC-like’ state. These include *Rps2* (> 99% of P5 cells but c. 28% of P6 cells), *Rps18* (> 99% of P5 cells but c. 60% of P6 cells), *Apoe* (c. 17% of P5 cells but c. 7% of P6 cells), and *Pou5f1* (c. 38% of P5 cells but c. 31% of P6 cells), and although these may guide further enquiry, we re-iterate that the purpose of this analysis was to provide an overview and that no formal statistical approach was undertaken to contrast the two populations.

***Re-assessing cell cycle phase using different gene marker sets***

Two of our main conclusions were that the majority of undifferentiated spermatogonia were predicted to be positioned in either the S or G_2_/M phases of the cell cycle (that is, actively replicating DNA), and that those cells in state 0B, more so than other transcriptomic states of undifferentiated spermatogonia, were disproportionately positioned in the G_2_/M phases (possibly to affect DNA repair). These conclusions were drawn on the basis of Seurat’s CellCycleScoring function, a marker gene-based means of cell cycle phase prediction which assigns either ‘G1’, ‘S’ or ‘G2’ phases to each cell. It does so by calculating an enrichment score for the S and G_2_/M phases using the average expression level of marker gene sets for both phases ^33^ (cells expressing neither S nor G_2_/M phase markers will have negative enrichment scores for both, and be assigned to G1 phase).

It follows that the predicted phase may be sensitive to the list of marker genes provided (although note that Seurat does not use the discrete G1/S/G2M classifications when regressing out the effect of cell cycle and so these cannot influence the resulting UMAP; rather, Seurat uses the quantitative ‘S phase’ and ‘G2M phase’ scores instead, as detailed at https://satijalab.org/seurat/articles/cell_cycle_vignette.html). For the purpose of creating the integrated atlas, we used a list of 67 experimentally-validated marker genes from a previous study ^34^ (hereafter, the ‘Dominguez’ set), as described in the main text. It is of interest to know to what extent our conclusions are influenced by this choice of gene list.

As such, to corroborate our findings, we re-predicted cell cycle phase for all cells in the SPG atlas using sets of S and G_2_/M marker genes made at varying levels of stringency. For this purpose, we used a curated longlist of 701 human S and G_2_/M-associated genes from a previous study ^35^ which monitored genome-wide expression in dermal fibroblasts as they synchronously entered the cell cycle from a quiescent (G_0_) state. For each gene on the longlist, there was an associated confidence score (from 1 to 13) reflecting the weight of evidence for its involvement in the cell cycle: one point for several lines of evidence from the study ^35^, whether it had been associated with a cell cycle-related phenotype in either human ^36^ or mouse ^37^, whether its knockdown led to a mitosis-related phenotype ^38^, and whether it had been identified in any of five previous human transcriptomic studies (^34,39–42^ [which includes the Dominguez set; **Supplementary Table 4**]). By filtering both on these confidence scores and on whether the cell cycle association was ‘known’ (widely accepted involvement with the cell cycle) or ‘putative’ (little or no direct evidence for involvement), we created 4 subsets of S and G_2_/M marker genes, of varying stringency, as follows:

(a) 701 genes, comprising 380 S genes and 321 G_2_/M genes,

(b) 496 ‘known’ genes, comprising 262 S genes and 234 G_2_/M genes,

(c) 387 ‘known’ genes with confidence score ≥ 5, comprising 210 S genes and 177 G_2_/M genes,

(d) 96 ‘known’ genes with confidence score ≥ 10, comprising 34 S genes and 62 G_2_/M genes.

As the enrichment of marker genes must by definition be relative, then the Seurat S and G2 phase scores can also be influenced by the composition of the cell population analysed. To minimise potential confounding effects or integration artefacts when predicting a cell’s phase of the cell cycle – using each of these four lists – we re-ran the Seurat workflow for each individual sample (that is, using unselected cells from each whole testis biopsy), rather than for the integrated dataset (that is, selected spermatogonia comprising the ‘SPG atlas’, where differences in cell cycle phase are unlikely to be as pronounced as those across the entire spermatogenic trajectory). The predicted cell cycle phase for each of the 4447 cells comprising the SPG atlas, both in the original atlas and using each of these four subsets, is given in **Supplementary Table 22**. From this table, we can make a number of observations. Firstly, the vast majority of phase predictions for the integrated atlas (which used the Dominguez set of markers) were identical to those made on a sample-by-sample basis with the same markers (3733 genes, 84% of the total), suggesting that although sample composition may influence the S and G2M scores, the impact was relatively minimal. Secondly, predictions from each of the four marker gene sets were largely concordant with those using the Dominguez set: for each of sets A, B, C, and D, there were 2605 (59%), 2638 (59%), 2586 (58%), and 3356 (75%) genes, respectively, with identical phase predictions to those made using Dominguez. Thirdly, and most importantly, irrespective of marker set, only a low proportion of cells were predicted to be in G_1_ phase (between 0 and 13% of the total) – that is, phase predictions discordant between datasets were largely because cells differed in being classified as S or G_2_ (**Supplementary Table 22**).

Overall, these results suggest that the choice of cell cycle phase marker genes does not alter the conclusion that undifferentiated spermatogonia are disproportionately found in the S and G_2_/M phases. It is also worth noting that irrespective of a given cell’s phase classification, many of the S and G_2_/M markers are detected in a relatively high proportion of cells in any given SPG atlas cluster anyway although in significantly more state 0B cells than those of either state 0 or state 0A/1 (**Supplementary Text Figure 1**). In addition, two G_2_/M markers functionally associated with DNA damage ^35^ are also seemingly ubiquitously expressed in state 0B: *XRCC6* and *HMGB2* (which are found in > 95% and >90% of state 0B cells, respectively, but 80-85% and 68-80% of cells in states 0 and 0A; **Supplementary Table 7**).

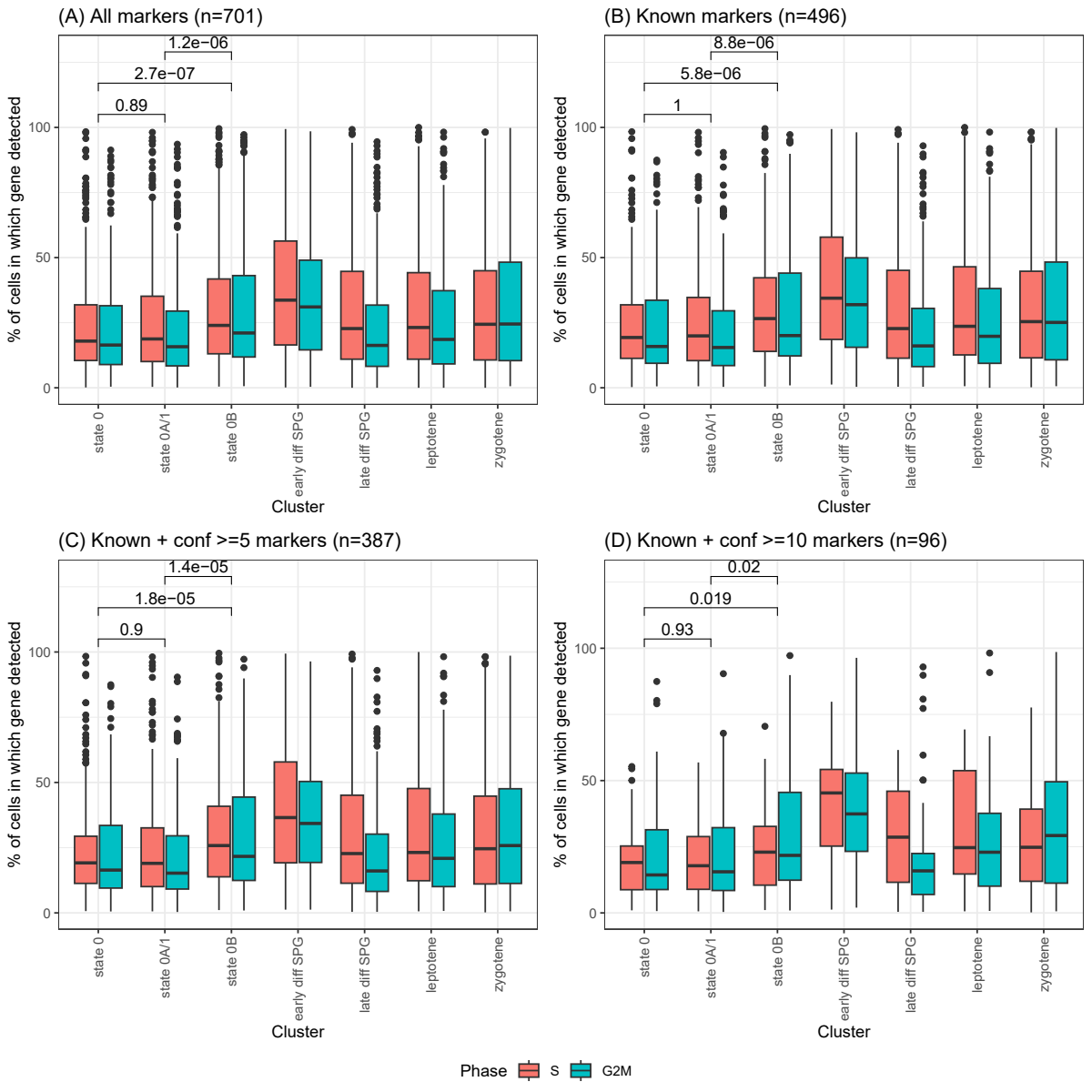

**Supplementary Text Figure 1.** Percentage of cells in each cluster of the SPG atlas (a proxy for expression level) in which four sets of S and G_2_/M-phase marker genes are detected (requiring ≥ 1 read per gene per cell).

The cell cycle phase marker gene lists are curated at varying levels of stringency: (a) 701 genes, (b) 496 genes, (c) 387 genes, and (d) 96 genes. Raw data for this figure are available in **Supplementary Table 7**. The p-values are those of a two-sided Wilcoxon rank sum test comparing the median percentage of cells per cluster in which either an S or G_2_/M marker gene is detected. It is apparent that genes characteristic of the S and G_2_/M phases are significantly more highly expressed in state 0B than in either states 0 or 0A/1, irrespective of the gene list used.

Nevertheless, the role of the G_2_/M phases in state 0B remains open to further, more detailed, enquiry. By assigning cells to one of only three predetermined phase categories, Seurat may not have correctly identified the actual phase *per se*. Other cell cycle prediction tools use different categories (for instance, Revelio has five ^43^ and CS1CC seven ^44^) although, as suggested by a recent benchmarking study ^33^, no tool yet shows obviously superior performance to any other; rather, for marker-gene-based methods as a whole, a key determinant of accuracy is how well suited the marker gene list is to the dataset being analysed. To that end, deriving a marker gene list specific to spermatogenesis (for a given species) could help shed further light on the nature of undifferentiated spermatogonia.

***The human state 0 transcriptomic program resembles, in part, that of mouse neuroblasts***

In the main text, we speculate that parallels exist between neural and spermatogonial stem cells as both could ‘rest’, pausing in the G_2_ phase of the cell cycle to affect DNA repair. To demonstrate that this is not a purely hypothetical possibility, here we show that neuroblasts (stem cell-like neural progenitors, the daughter cells of transit-amplified neural stem cells) can be projected onto the human SPG atlas, and that when they do they resemble cells of the ‘state 0’ cluster. This observation opens the possibility for further exploration of the parallels between the two systems and to what extent they resemble each other functionally. This could potentially be a fruitful line of enquiry as there are, in general, a relatively high number of cellular and molecular similarities between neurons and sperm ^45^.^[3]*^

For this analysis, we obtained a scRNA-seq dataset (SRA accession SRS6956708) from a 5 week old mouse, used in a previous study to characterize neurogenesis ^46^. More specifically, this study showed that the transcriptomic profile of striatal astrocytes (a support cell found only in a specific brain region) resembled that of a neural stem cell, as in conditions approximating the aftermath of a stroke (deletion of the Notch-mediating transcription factor *Rbpj*, which mimics the impact of reduced Notch signaling) they could generate new neurons. We processed the SRS6956708 dataset using the same workflow and parameters as the mouse SSC datasets (see Materials and Methods), then projected it onto the SPG atlas UMAP (**Figure 1B**) using the same two-step projection approach, i.e. by first projecting cells onto the whole-testes UMAP (**Figure 1A**) and retaining only those which map to the germline (n = 361 cells). We can see that cells from this dataset project onto the SPG atlas only on the right-hand side of the ring: primarily onto the state 0 (n = 288 cells) and early diff SPG (n = 48 cells) clusters (**Supplementary Text Figure 2**). To interpret this result, we need to consider what, specifically, the SRS6956708 dataset comprises: these are FACS-sorted astrocytes and their neurogenic progeny, isolated from the striatum of a mouse homozgyous for *Rbpj* knockout. Accordingly, the dataset captures a neurogenic trajectory, from astrocytes to transit-amplifying cells to neuroblasts, alongside additional support cells (oligodendrocytes), as illustrated in their ‘figure 1 – figure supplement 1’: https://www.ncbi.nlm.nih.gov/pmc/articles/PMC7440914/figure/fig1s1/. To confirm which neurogenic cell type projected onto each of the state 0 and early diff SPG clusters, we projected each set of cells back onto the SRS6956708 UMAP (**Supplementary Text Figures 3 and 4**, respectively), noting that this UMAP – although created using a different method to ^46^, namely the Kallisto/Bustools/Seurat workflow – is in essence identical to their ‘figure supplement 1’. We find that cells which projected onto the ‘early diff SPG’ cluster originated from the ‘transit-amplifying’ cluster (**Supplementary Text Figure 3**) – consistent with the fact that both are transient, not fully differentiated, populations – and that cells which projected onto the ‘state 0’ cluster originated from the ‘neuroblast’ cluster (**Supplementary Text Figure 4**).

What is striking about this observation is that the neurogenic and spermatogenic trajectories, ostensibly originating in ‘astrocyte’ and ‘state 0’ populations, respectively, do not run in parallel with each other. This suggests that there are specific transcriptional similarities between the ‘state 0’ and ‘neuroblast’ populations that the projection algorithm is capturing instead; these may provide a fruitful direction for future research. Notably, in vertebrates, neuroblasts are considered ‘young neurons’ that exist in a state of transient mitotic quiescence prior to their terminal differentiation (upon which they become irreversibly ‘locked in’ to G_0_ phase) ^47^. It is conceivable, therefore, that the mechanisms by which neuroblasts enter, exit, or are ‘paused’ in the cell cycle may shed further light on how undifferentiated spermatogonia behave.

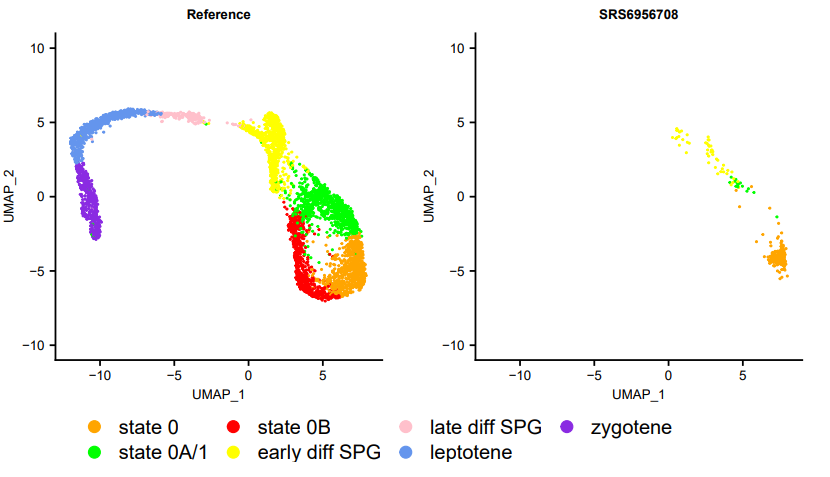

**Supplementary Text Figure 2.** Projection of a mouse neural dataset (n=361 cells) onto the SPG atlas (the ‘reference’ UMAP).

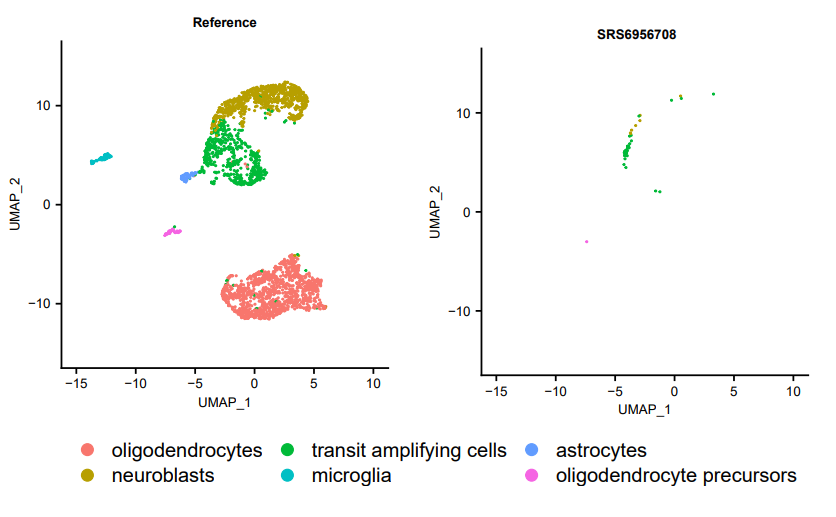

**Supplementary Text Figure 3.** Re-projection of ‘early diff SPG’-resembling mouse neural cells from sample SRS6956708 onto the ‘reference’ UMAP, i.e. the full set of cells from sample SRS6956708.

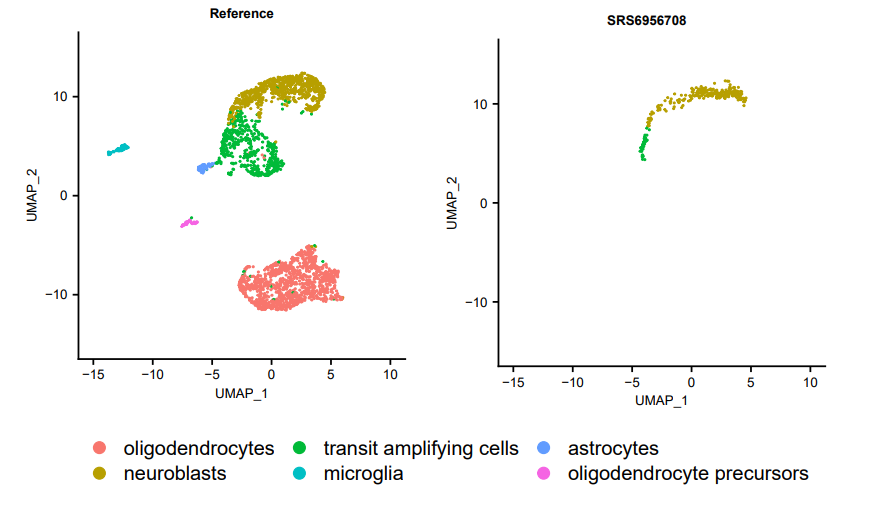

**Supplementary Text Figure 4.** Re-projection of ‘state 0’-resembling mouse neural cells from sample SRS6956708 onto the ‘reference’ UMAP, i.e. the full set of cells from sample SRS6956708.

***Projection of a time course of mouse germ cells onto the SPG atlas***

In **Figure 4**, we show the projection of selected mouse germ cell samples onto the SPG atlas, representing a time course from early development to adulthood. These samples were selected to emphasise early time points in particular (in which heterogeneity of the undifferentiated spermatogonial compartment was more pronounced) and one representative example of an adult (4 month) mouse. Below, in the composite **Supplementary Text Figure 5**, we show the projection of each of the 37 samples. Details of the samples included in this analysis are given in **Supplementary Table 2**, with the relative abundance of each cluster across the full set of 37 samples shown in **Figure 2B**. The number of cells projected onto each cluster, for a range of minimum projection scores (0 to 0.8 at 0.2 intervals), are given in **Supplementary Table 17**, with the following figures plotted with no minimum threshold required.

Note that as these samples have been drawn from multiple studies and because we have implemented a uniform set of reasonably conservative data-quality filters, they may vary in terms of their apparent quality (that is, the number and type of cells represented). One of the conclusions of our study is that there is no compelling evidence that human-like states 0 or 0B are present in adult murine spermatogonia, although we cannot ignore the fact that a small number of (ostensibly lower-quality) adult samples do appear to show a small number of projected ‘state 0’ cells (but not state 0B). However, we disregard these observations as artefactual on account of the fact that these are not consistently repeated findings (unlike, for instance, the presence of human-like state 0A, which is apparent in every projection), that they cannot be replicated in rats (**Figure 5**), and that these projections to ‘state 0’ only occur when no minimum confidence threshold is required; that is, when the projection of any cell is forced.

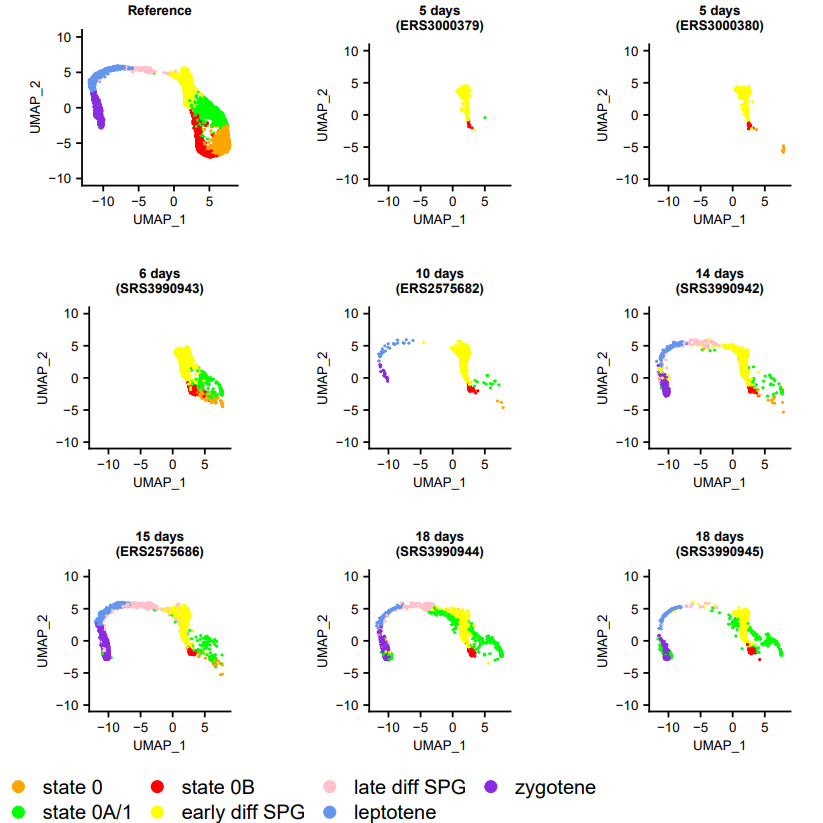

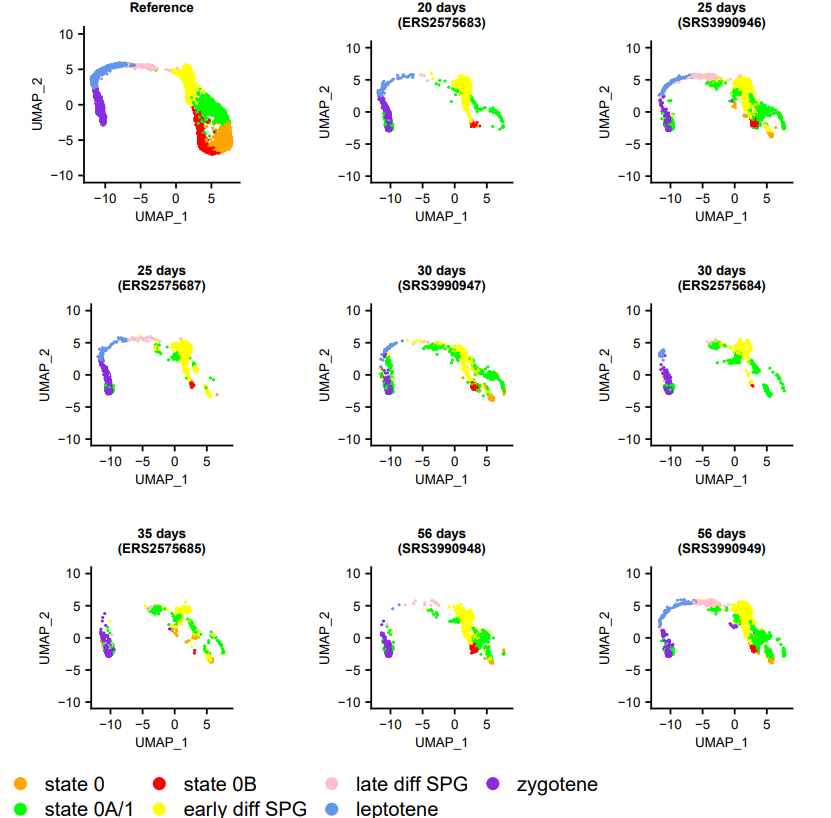

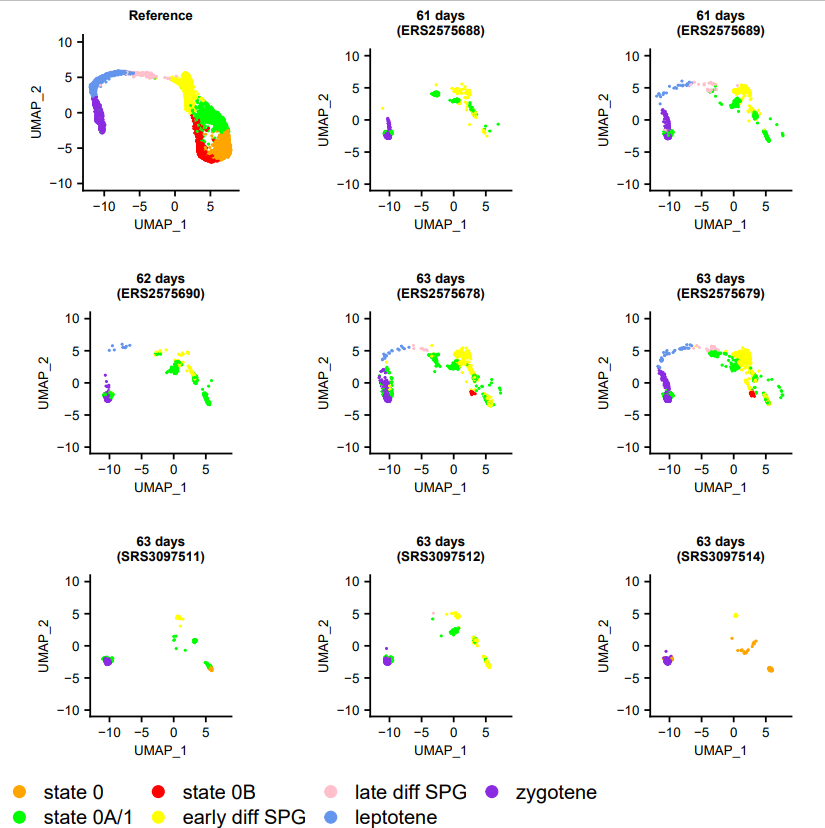

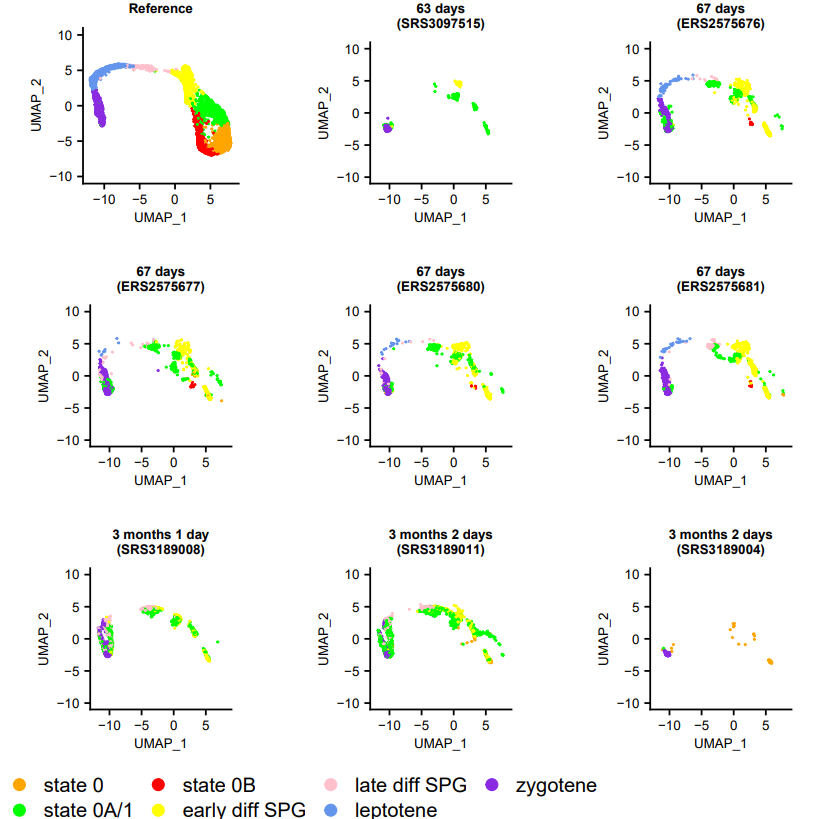

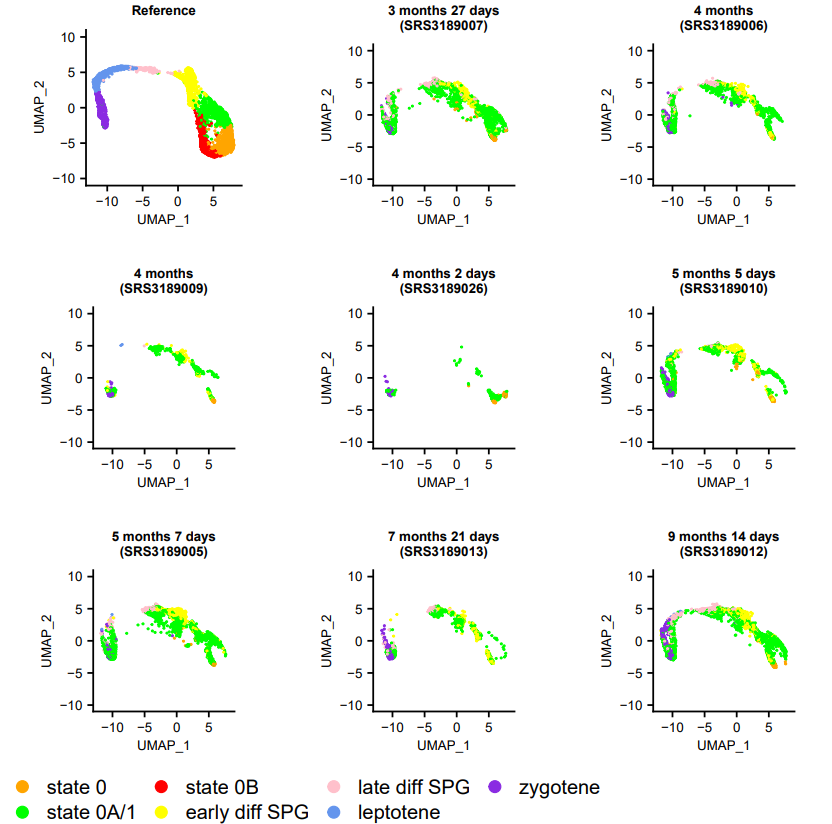

**Supplementary Text Figure 5.** Projection of mouse germ cells onto the SPG atlas, using 37 representing a time course from early development to adulthood.

***Projection of a time course of human germ cells onto the SPG atlas***

In the main text, we assessed whether each of the undifferentiated cell states were present throughout the human life cycle, by conservative projection of a time course of 62 samples, inclusive of the 34 samples used to construct the atlas, and incorporating foetal (8-23 week), pre- and peri-pubertal timepoints. Below, in the composite **Supplementary Text Figure 6**, we show the projection of each of these samples onto the SPG atlas. Details of the samples included in this analysis are given in **Supplementary Table 2**, with the relative abundance of each cluster across the full set of 62 samples shown in **Figure 2A**. The number of cells projected onto each cluster, for a range of minimum projection scores (0 to 0.8 at 0.2 intervals), are given in **Supplementary Table 15**, with the following figures plotted with no minimum threshold required. Owing to the conservative thresholds employed to integrate the data, individual samples may contain only a small number of cells; as such, the projections summarised in **Figure 2A** use aggregated data across age categories. Note also that this figure includes a projection to an additional sample from ^18^ by way of validation: a 6-week-old embryo (run accession HRR131888), the cells of which as expected do not project onto the SPG atlas (because humans do not undergo sex differentiation until approximately the seventh week of gestation) ^48^.

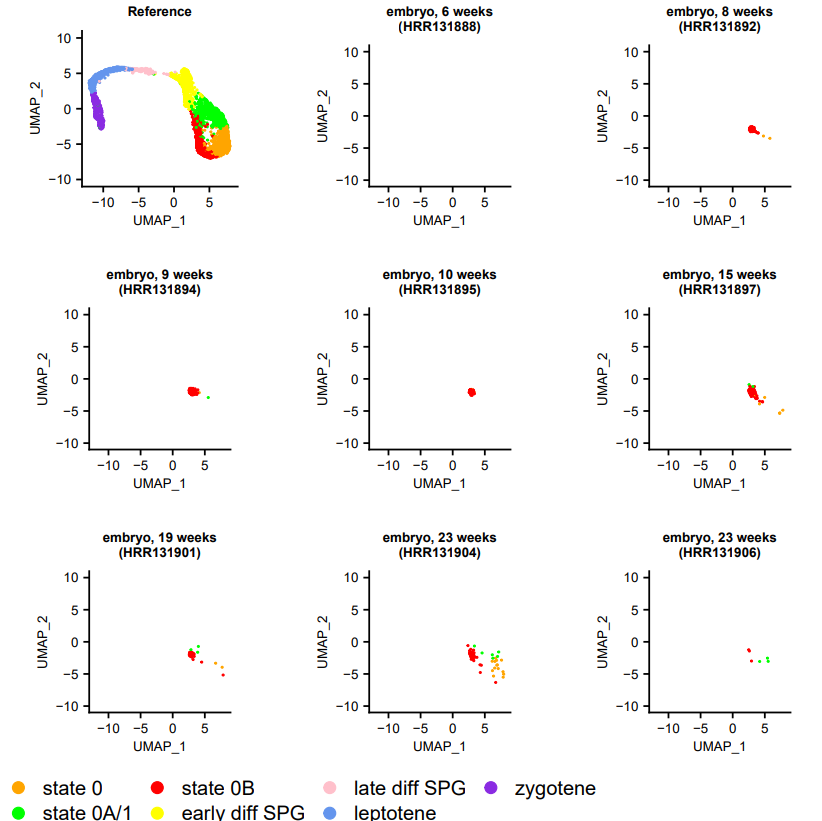

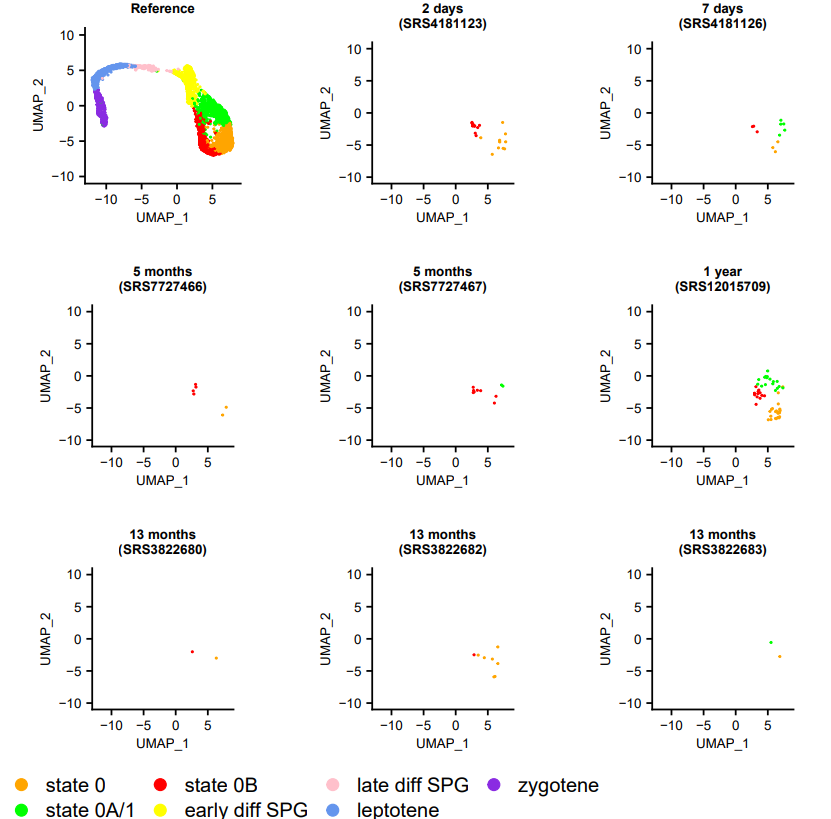

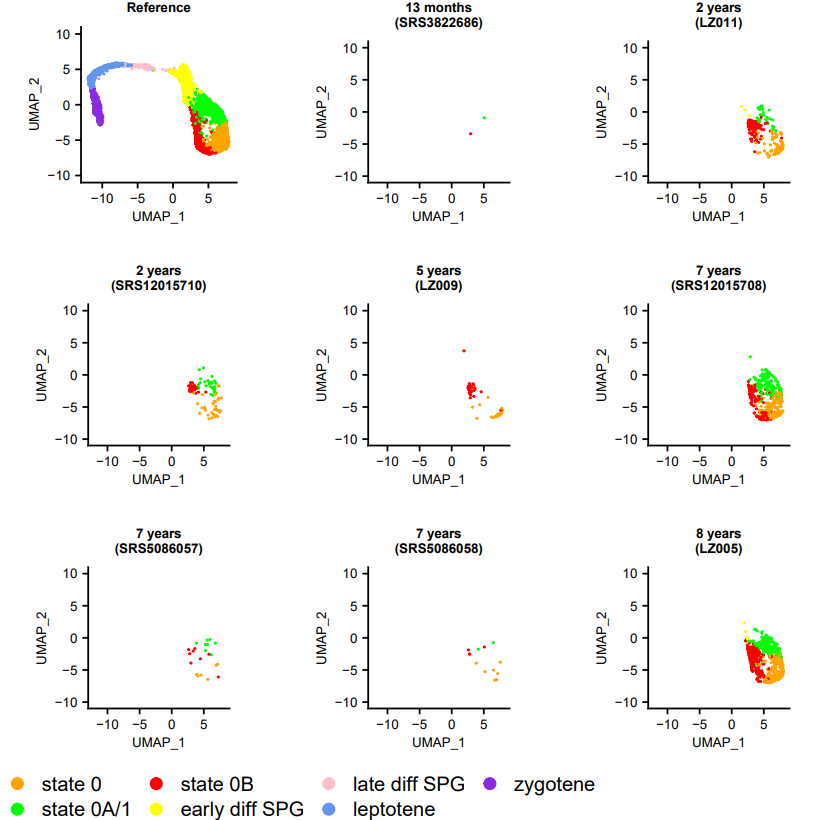

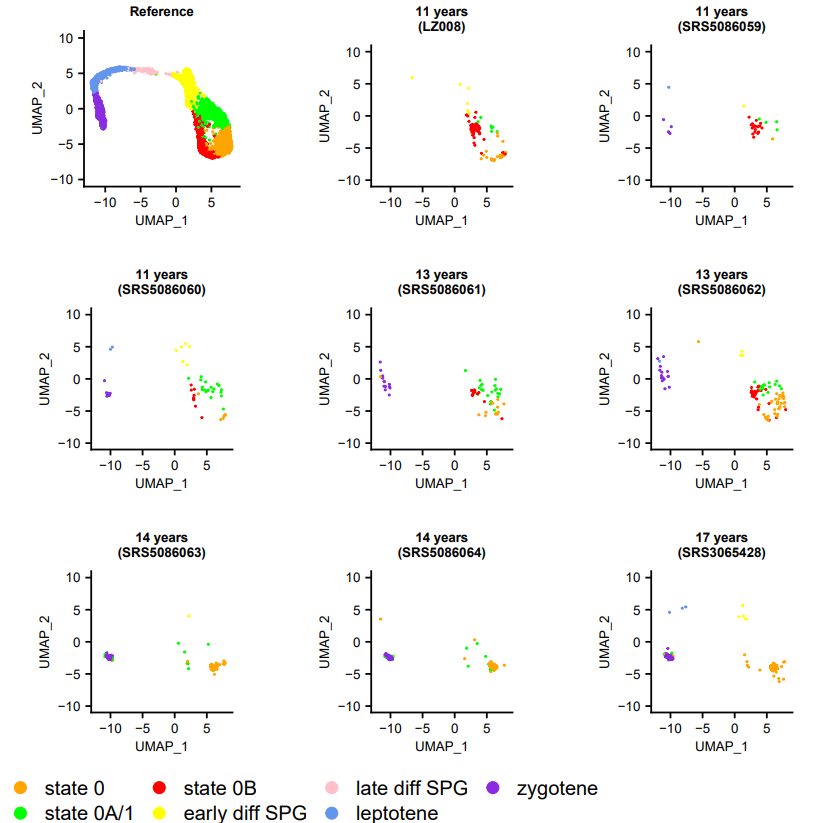

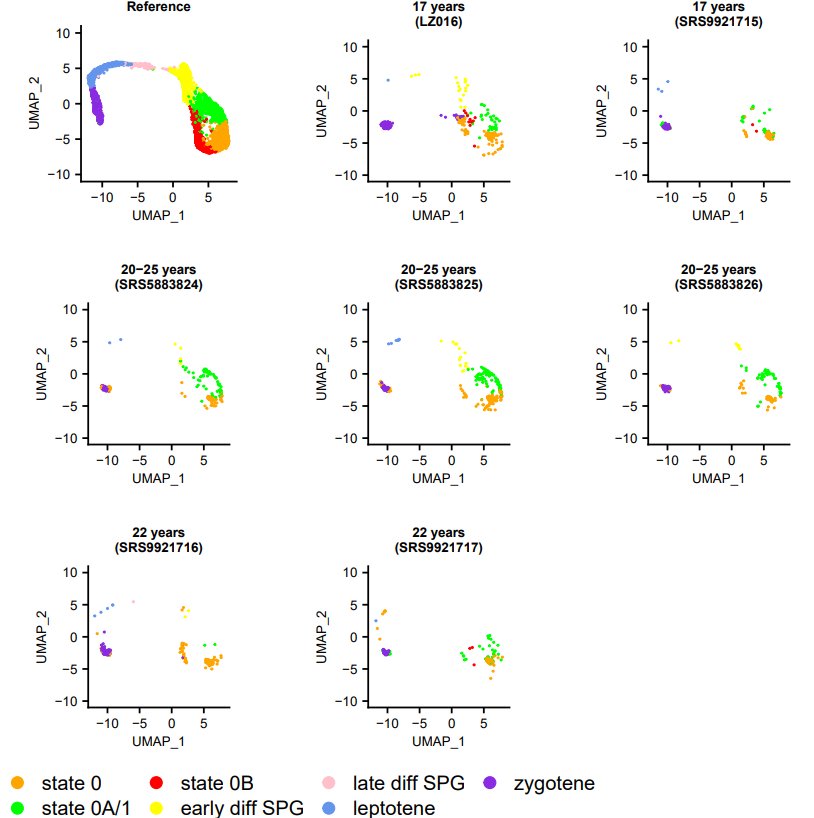

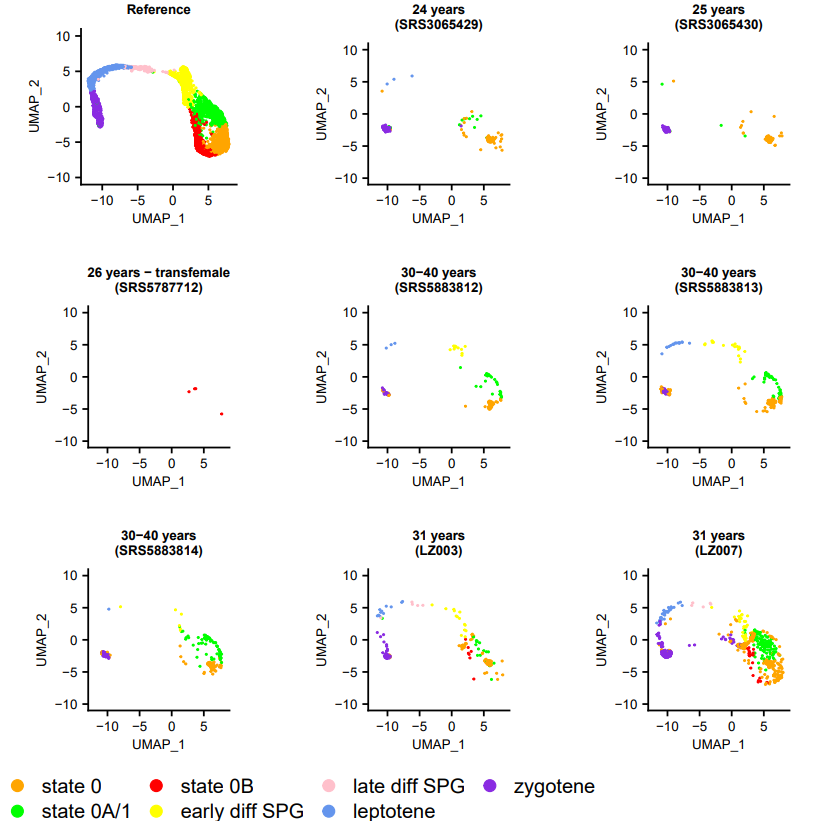

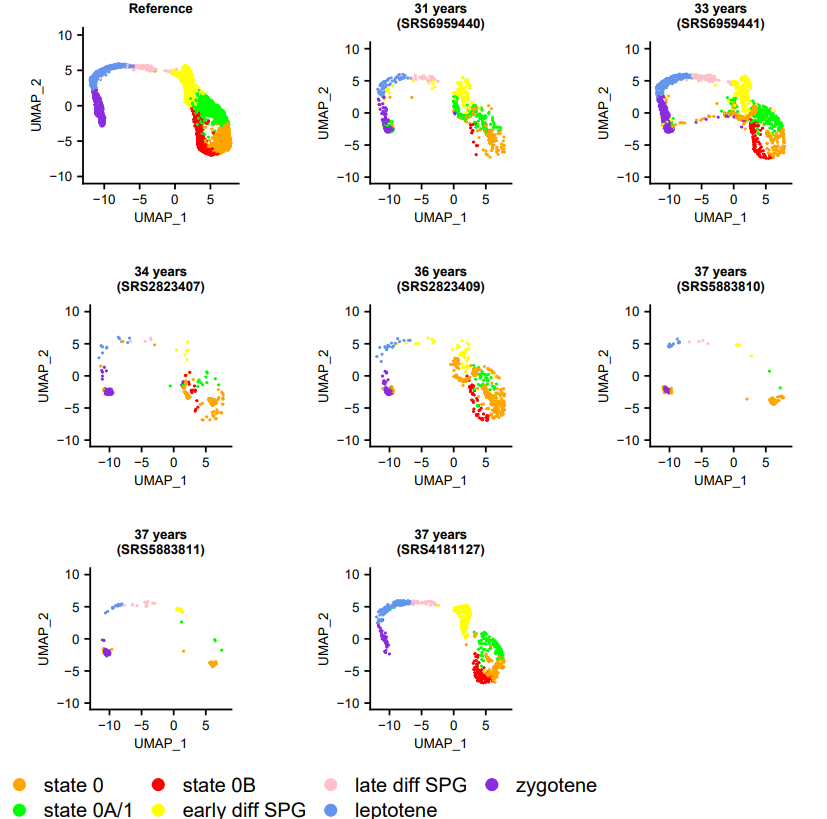

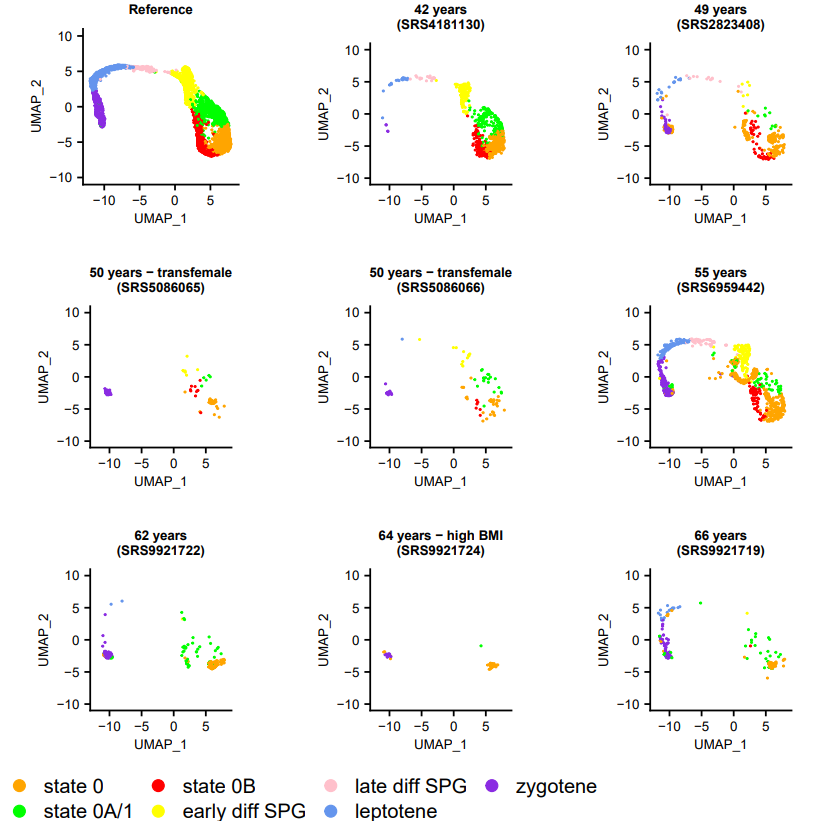

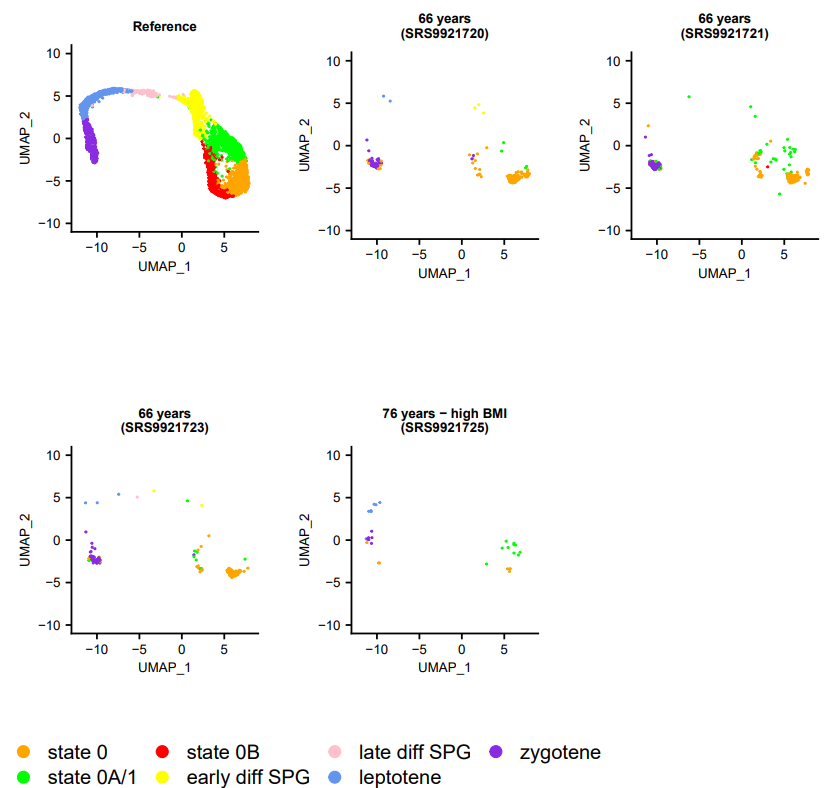

**Supplementary Text Figure 6.** Projection of human germ cells onto the SPG atlas, using 62 representing a time course from early development to adulthood.

1. *	When clustering the SPG atlas at resolution 1.1, *NANOS2*, *CCDC106*, *RPL3*, *SCAMP2*, *XAB2*, *SGF29*, and *TMEM39B* are differentially expressed only in ‘state 0B’. Although two subsets of state 0B form at resolution 1.2 and higher (which we refer to as ‘state 0/0B’ and ‘state 0B/1’; **Supplementary Figure 8**), it is worth noting that each of these seven genes are also differentially expressed in both of them (**Supplementary Table 8**) and so the following observations apply irrespective of resolution. Other ‘state 0B’ genes referred to in the main text that are also DE in both subsets are the cell-cycle-associated *MDH1* and the DNA-damage-associated *PSTK* and *TMEM39B* (**Supplementary Table 8**).
2. *	Note that orthology relationships were obtained from Ensembl BioMart v104 ^19^, and that on the basis of their gene models, human *SETD1A* (ENSG00000099381) is not considered an orthologue of mouse *Setd1a* (ENSMUSG00000042308).
3. *	Also supporting a link between neurons and sperm is the incidental observation that in our data the most highly	enriched KEGG pathways for each cluster in the SPG atlas – relative to the state 0 ‘apex’ – are ‘Alzheimer’s	disease’, ‘Huntington’s disease’ and ‘Parkinson’s disease’, signalling pathways the dysregulation of which are	associated with neurological disorders (**Supplementary Table 12**).
